# The intersection of endocrine signaling and neuroimmune communication regulates neonatal nociception

**DOI:** 10.1101/2024.07.26.605393

**Authors:** Adewale O. Fadaka, Adam J. Dourson, Megan C. Hofmann, Prakriti Gupta, Namrata G.R. Raut, Michael P. Jankowski

## Abstract

Neonatal pain is a significant clinical issue but the mechanisms by which pain is produced early in life are poorly understood. Our recent work has linked the transcription factor serum response factor downstream of local growth hormone (GH) signaling to incision-related hypersensitivity in neonates. However, it remains unclear if similar mechanisms contribute to inflammatory pain in neonates. We found that local GH treatment inhibited neonatal inflammatory myalgia but appeared to do so through a unique signal transducer and activator of transcription (STAT) dependent pathway within sensory neurons. The STAT1 transcription factor appeared to regulate peripheral inflammation itself by modulation of monocyte chemoattractant protein 1 (MCP1) release from sensory neurons. Data suggests that STAT1 upregulation, downstream of GH signaling, contributes to neonatal nociception during muscle inflammation through a novel neuroimmune loop involving cytokine release from primary afferents. Results could uncover new ways to treat muscle pain and inflammation in neonates.

**Graphical Abstract:** 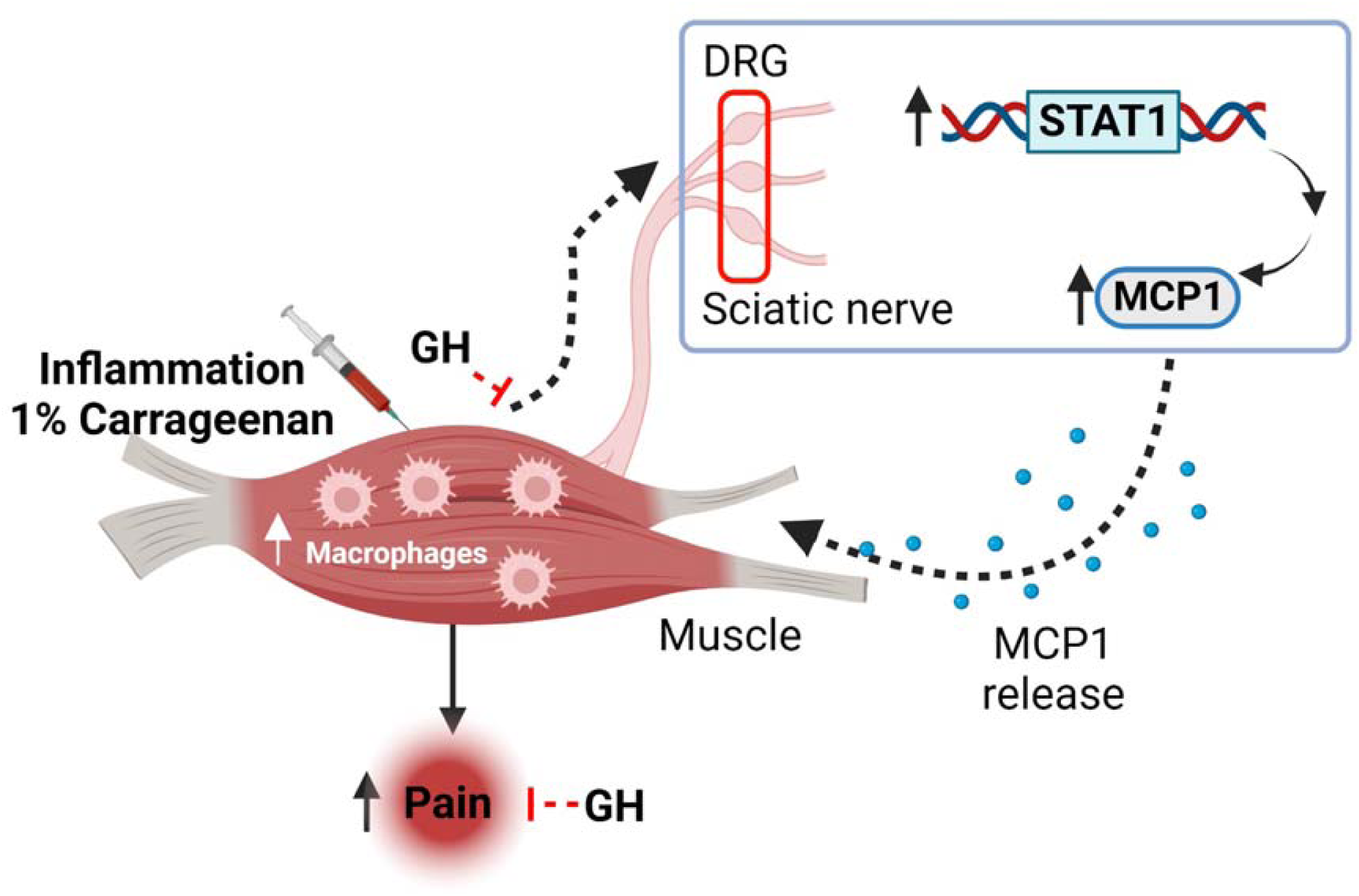

## Introduction

Nociceptive signals in neonates are thought to be transduced from the periphery to the central nervous system (CNS) distinctly from adults [1] . Further, painful events occurring during early neurodevelopmental stages when the nervous system is immature, result in [2, 3] prolonged modifications in pain processing [4] and even neurodevelopmental impairments [5]. This highlights the importance of understanding unique mechanisms of nociceptive processing in neonates.

Growth hormone (GH) exerts its function by binding to growth hormone receptor (GHR) [6]. Macrophages express GHR on their surface and are directly responsive to GH which in turn regulates cytokine production [6, 7]. GH deficiency in neonates alone induces behavioral and afferent hypersensitivity to peripheral stimuli while deletion of GHr from primary afferents similarly induces peripheral hypersensitivity [8, 9]. GH has also been shown to regulate afferent sensitization and responsiveness to peripheral injury during incision injury through a unique interaction between immune cells and sensory neurons which can influence nociceptive responses later in life [9, 10]. This suggests a critical involvement of GH in neonatal nociception and nociceptive priming.

GH exerts different pleiotropic effects through unique intracellular signaling pathways. GH mainly activates the Janus kinase/signal transducer and activator of transcription (JAK/STAT) signaling pathways [11] and both JAKs and STATs transduce signals initiated by many growth factors and cytokines [12]. JAKs can also interact with transcription factors (TFs) to modulate gene expression [13]. Activation/ suppression of many transcription factors are associated with the development of hyperalgesia after injury [14]. One group of TFs induced particularly in sensory neurons after inflammatory injury [15, 16] and contributing to inflammatory pain development is the STAT family [17]. Persistent activation of STATs is known to promote chronic inflammation [18]. STAT proteins share the same structural motif but are often involved in different biological processes. *STAT1* as a prototypical member has been reported to be activated by cytokines and growth factors including GH and insulin [19, 20]. Mice with *STAT1* knockdown exhibit selective signaling defects in their response to both Type-1 and Type-II interferons [21] but respond normally to several other cytokines that activate *STAT1*. However, *STAT1* is far less studied when compared with *STAT3* and *STAT5,* thus its role in neonatal inflammatory pain deserves closer attention.

Macrophages are central to pathogen recognition and responses to injury for purposes of tissue repair [22, 23]. These cells can be primed and modulated by GH [24] resulting in the controlled release of proinflammatory cytokines and improved macrophage function by increasing superoxide and TNFα release [25, 26]. Our recent work has linked the transcription factor SRF and local GH signaling to incision-related hypersensitivity in neonates through a coordinated interaction with infiltrating macrophages [27]. However, it is not known if similar transcriptional mechanisms within sensory neurons contribute to inflammatory pain in neonates. Therefore, the current study investigated if distinct neuronal transcription factors or local GH may modulate nociceptive behaviors in neonatal mice following muscle inflammation. Here, we uncovered a new role for peripheral GH in neonatal inflammatory myalgia. We found that GH modulated inflammatory muscle pain in neonates through neuronal expression of STAT1, but the mechanism by which this pathway exerted its influence on neonatal pain was through its modulation of non-peptidergic, neurogenic inflammation through the release of the cytokine monocyte chemoattractant protein 1 (MCP1, aka. CCL2) from sensory neurons into the injured muscle.

## Results

### Local GH modulates pain-related behaviors after muscle inflammation

To confirm whether muscle inflammation contributes to pain-related behaviors in neonatal mice, we used 1% carrageenan injection into the hind paw muscles to induce inflammation at postnatal day 7 (P7). Muscle inflammation increased spontaneous paw guarding scores, and lowered paw withdrawal thresholds at day 1 post inflammation compared to BL and naïve/sham mice (Figure 1). Similar results were found after cutaneous inflammation (Figure S1). Minimal effects were observed in the contralateral paw after muscle inflammation in regard to guarding behaviors. However, there was a slight reduction in mechanical withdrawal thresholds (Figure S1). These data suggested that the muscle inflammation induced pain-related behaviors in neonatal mice are similar to what has been observed in adults [28, 29].

**Figure 1:**
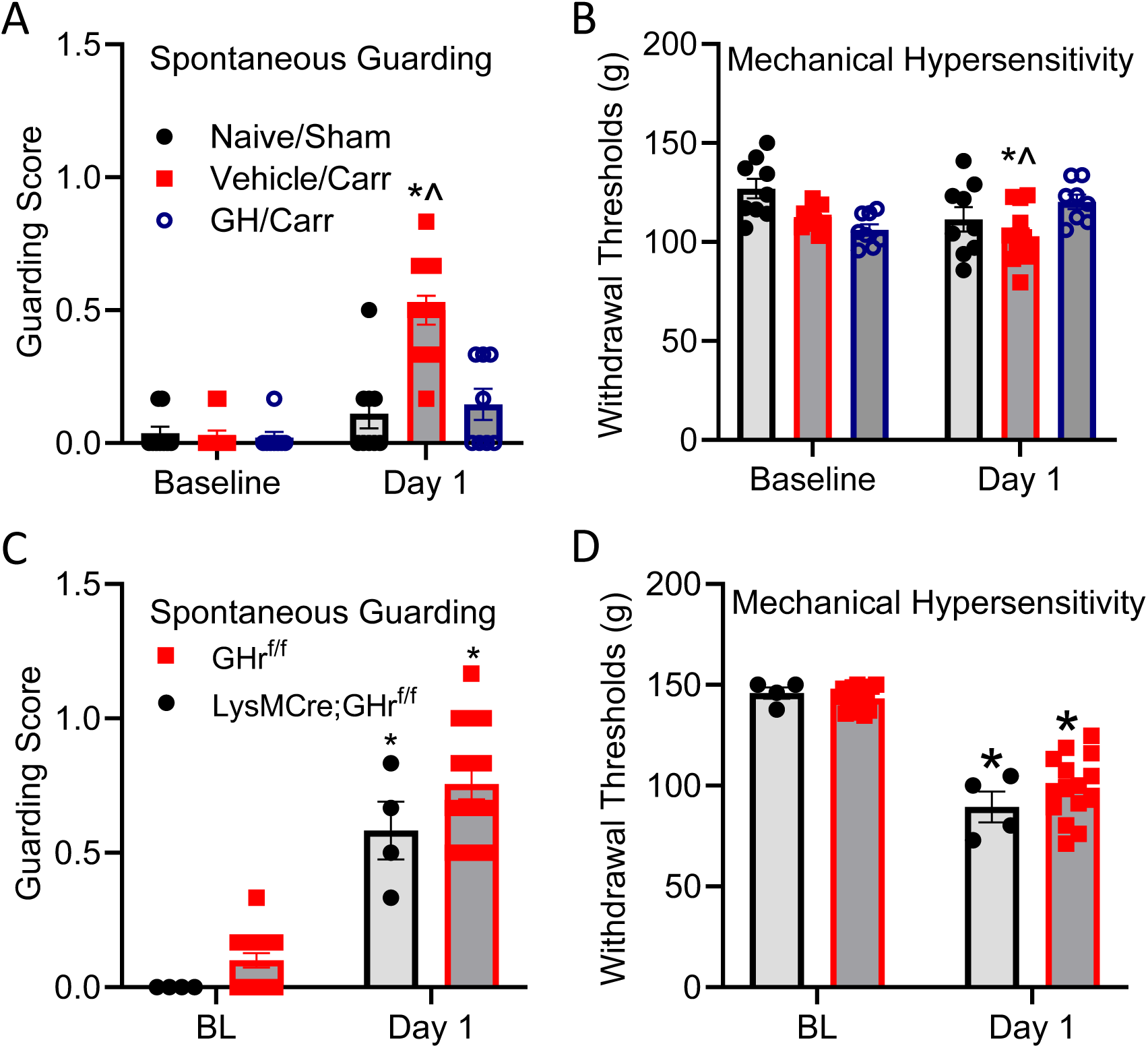
GH modulates nociceptive behaviors after muscle inflammation in neonates. **A.** One day after carrageenan induced muscle inflammation in neonatal mice at P7, animals display increased paw guarding behaviors. This is inhibited by local GH treatment. (*p<0.05 vs. baseline (BL) and ^p<0.05 vs. Carrageenan+Vehicle (Carr/Veh)), **B.** Muscle inflammation decreases mechanical withdraw thresholds at 1 day (*p<0.05 vs. BL and ^p<0.05 vs. Carr/Veh), and this is also blocked by GH injection into the muscles of inflamed mice. **C:** Knockout of GHr from macrophages (LysMCre;GHr^f/f^) does not alter the increased spontaneous paw guarding observed 1d after muscle inflammation. **D:** No changes in mechanical hypersensitivity are found in neonatal mice with GHr knocked out in macrophages compared to littermate controls with muscle inflammation. Statistics: Behavioral results are expressed as mean ± SEM (n = 4-12). Two-way RM ANOVA followed by Tukey’s post hoc multiple comparison test.

We then wanted to confirm that a local GH treatment could reverse muscle inflammatory myalgia in neonates similar to our recent reports after muscle incision [27]. Behaviorally, the local GH intervention in the muscles significantly decreased spontaneous paw guarding (Figure 1A) and blocked the reduction in mechanical withdrawal thresholds (Figure 1B) induced by muscle inflammation with 1% carrageenan. Although GH treatment also restored contralateral mechanical hypersensitivity in mice with muscle inflammation, GH treatment had no effects on body weight (Figure S1). This indicated that GH has a similar anti-nociceptive effect in neonates with muscle inflammation as has been found after incision injury at day 1. We therefore wanted to confirm if this was due to a similar sequestering of GH by infiltrating macrophages. In mice with the GH receptor (GHr) knocked out in macrophages (LysMCre;GHr^f/f^), we surprisingly found that carrageenan induced hypersensitivity was not blocked in these animals compared to littermate controls (Figure 1C-D). This suggested a unique mechanism by which GH could inhibit pain-related behaviors in neonates after an inflammatory insult.

### GH suppresses sensory neuron sensitization induced by muscle inflammation in neonatal mice

As muscle inflammation induced behavioral hypersensitivity in neonates but GH appeared to not affect these behaviors via its actions on infiltrating macrophages under these specific injury conditions, we wanted to determine if muscle inflammation induced sensory neuron sensitization could be inhibited by local GH treatment. We therefore employed a novel *ex vivo* hind paw muscle, tibial nerve, DRG, spinal cord calcium imaging preparation using genetically encoded Ca^2+^ indicators expressed in sensory neurons (PirtCre;GCaMP6) in neonatal mice subjected to muscle inflammation with or without local GH intervention (Figure 2A). Muscle afferents innervating the flexor digitorum brevis muscles showed a variety of responses to mechanical, thermal (hot and/or cold) and chemical (Low and/or high metabolite mixtures containing ATP, lactate and protons) stimuli (Figure 2B, C). Primary afferents in mice with muscle inflammation showed a reduction in mechanical thresholds when compared to naïve mice. Treatment of inflamed mice with the GH intervention reversed mechanical hypersensitivity in the group III/IV muscle afferents (Figure 2D). Although no changes in the total numbers of mechanically sensitive afferents were found in any of the three conditions, we did observe a significant increase in the number of cells responding to heat stimuli as well as an increase in cells responding to both metabolite mixtures. Local GH treatment in mice with muscle inflammation however showed proportions similar to naive (Figure 2E). There was no significant difference in the mean intensity of the response to mechanical and chemical (high, low and H/L) stimulation of the receptive fields in the muscles between the groups (Figure 2F). No changes in DRG neuron responsiveness were observed over increasing mechanical forces either (Figure S2). For heat response intensity however, mice with muscle inflammation show an increase in mean florescence intensity compared to naïve (p=0.0305) but this is not statistically different in inflamed mice treated with GH (p=0.1253) (Figure 2F). This data suggests that GH may modulate inflammatory myalgia in neonates at the level of the primary afferent.

**Figure 2:**
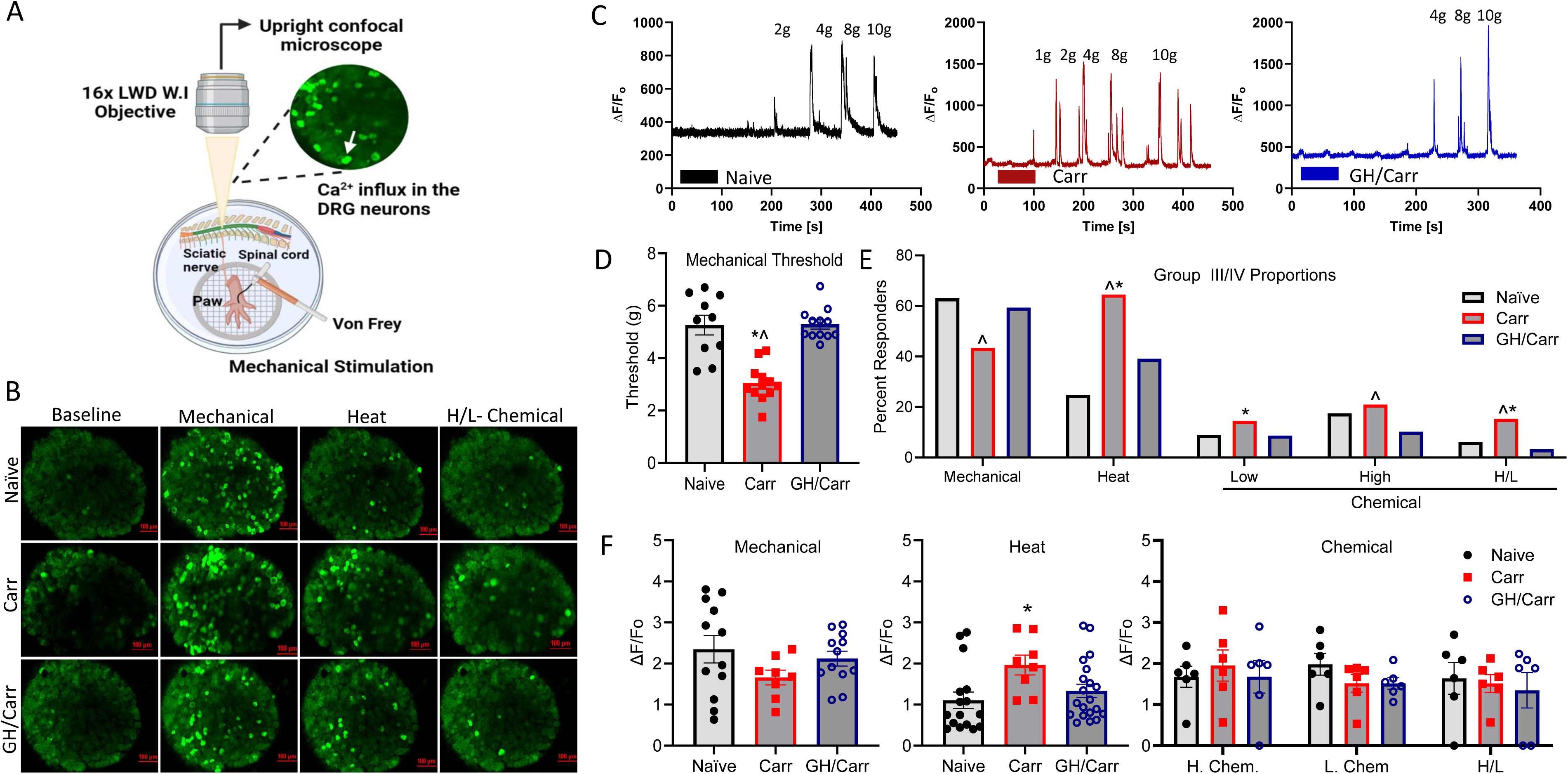
*Ex vivo* calcium imaging of the DRGs in neonates with hindpaw muscle inflammation with or without GH treatment. **A.** Schematic representation of the calcium imaging *ex vivo* hind paw muscle/tibial nerve/DRG/ spinal cord preparation and data recording in DRG neurons expressing GCaMP6f. **B.** Representative images of DRGs from *ex vivo* preparations in PirtCre;GCaMP6f mice before (BL) and after different stimulations (mechanical, heat, and H/L-chemical) of the muscle receptive fields. **C.** Responses of DRG neurons of mice expressing GCaMP6f in sensory neurons following exposure to mechanical stimuli in the three groups (Naïve, Carrageenan (Carr) and GH+ carrageenan treatment (GH/Carr)). **D.** Mechanical thresholds of sensory neurons are reduced after muscle inflammation when compared with naïve (*p<0.0001 vs. naïve) but this is blocked in mice with the GH intervention (^p<0.0001 vs. GH+Carr). **E.** There are no differences in the number of mechanically sensitive cells after muscle inflammation with or without GH intervention compared to naïve animals. However, there is a significant increase in the number of heat sensitive cells after muscle inflammation compared to naïve mice (*p= 0.0117 vs naïve). This effect is not observed in mice with carrageenan injection into the muscles plus the GH intervention (p= 0.0846 vs GH/Carr). Chemically sensitive cells were categorized into cells that respond to a high concentration metabolite mixture (H. Chem: 50mM lactate, 5μM ATP, pH 6.6), a low concentration metabolite mixture (L. Chem: 15mM lactate, 1μM ATP, pH 7.0), and cells that respond to both high and low metabolite mixtures (H/L). There was an increase in cells that responded to both mixtures after muscle inflammation compared with naïve. This effect was also significantly inhibited by the GH intervention (^p= 0.0254 vs GH/Carr and p=0.053 vs naive). **F.** There is no significant difference in the mean florescence intensity of sensory neurons to mechanical stimulation of the RFs after muscle inflammation when compared to naïve (p=0.1914 vs. naïve) or GH intervention groups (p=0.4636 vs. GH/Carr). There is a significant increase in calcium florescence in sensory neurons to heat stimuli after muscle inflammation compared to naïve groups (*p= 0.0305 vs naïve). This was partially (non-significantly) reduced in inflamed mice with a GH intervention (p=0.1253 vs. GH/Carr). For metabolite florescence intensity in sensory neurons, the observed difference after muscle inflammation is significant only when compared to the GH+carrageenan treated group (*p= 0.0224 vs GH/Carr) but not with naïve group (p=0.0892 vs. naives). Stats: Mechanical threshold, *p=0.0001 vs. naïve; ^p<0.0001 vs. GH+1%Carr group. One-way ANOVA, Tukey’s post hoc. Cell numbers= 297 (naïve), 174 (Carr), and 447 (GH+Carr) from 3 mice each. Mean ± SEM. Cell responder proportions, mechanical (naive vs Carr.; χ² = 2.135, 1 df, p=0.1439; naive vs GH/Carr.; χ²= 0.8424, 1 df, p=0.3587; GH/Carr vs Carr.; χ² = 5.176, 1 df, ^p=0.0229), heat (naive vs Carr.; χ² = 32.73, 1 df, *p<0.0001; naive vs GH/Carr.; χ²= 5.238, 1df, *p=0.0221; GH/Carr vs Carr.; χ² = 12.65, 1 df, ^p=0.0004), and chemical [low (naive vs Carr.; χ² = 1.841, 1 df, *p=0.0.1748; naive vs GH/Carr.; χ²= 0.000, 1 df, p>0.9999; GH/Carr vs Carr.; χ² = 1.841, 1 df, p=0.1748), high (naive vs Carr.; χ² = 0.299, 1 df, p=0.5844; naive vs GH/Carr.; χ²= 2.101, 1 df, p = 0.1472; GH/Carr vs Carr.; χ² = 3.929, 1 df, ^p=0.0475) or both (naive vs Carr.; χ² = 0.4.315, 1 df, *p=0.0378; naive vs GH/Carr.; χ²= 1.048, 1 df, p = 0.3061; GH/Carr vs Carr.; χ² = 8.800, 1 df, ^p=0.003)] responsive muscle afferents are altered by inflammation and rescued by GH intervention.

### GH regulates muscle inflammation

In the process of confirming a role for GH in muscle inflammation related anti-nociception, we noticed that there appeared to be local effects of GH treatment during this specific injury that was not observed in our previous report assessing incision injury [1]. As expected, muscle inflammation with 1% carrageenan significantly induced paw edema (i.e. increased paw volume) in neonates compared to naïve mice. However, the effects of muscle inflammation were partially inhibited following GH treatment. (Figure 3A). No changes in paw volume were observed in the contralateral paw (Figure S3).

**Figure 3:**
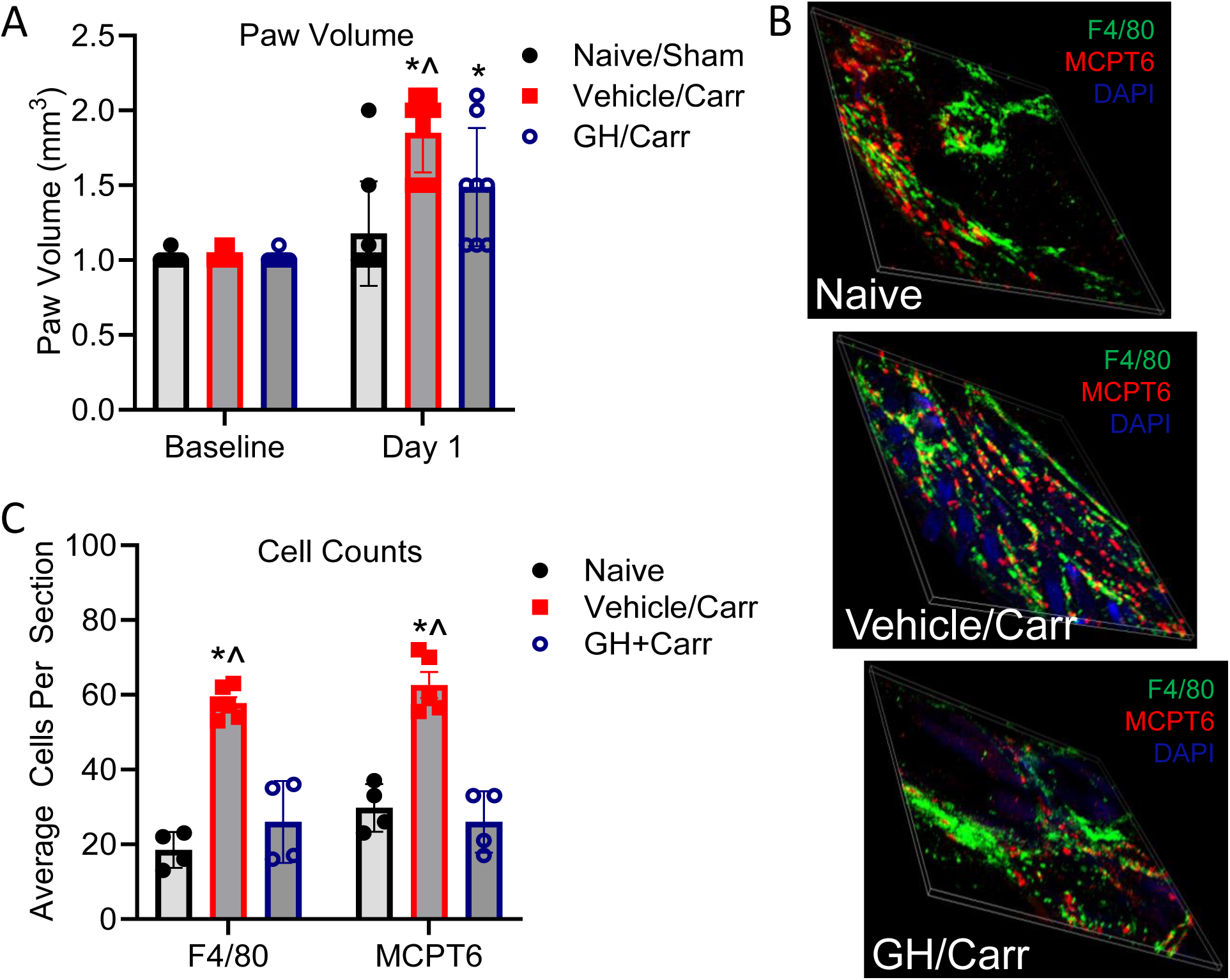
GH modulates paw edema and immune cell infiltration in the muscles of neonates. **A.** 1% carrageenan injection into the hind paw muscles of P7 neonates increases paw volume but this is reduced by local GH treatment (*^p<0.05 vs BL and Carr/Veh). **B.** Volume imaging of immune cells in the muscles after inflammation or inflammation with GH intervention compared to naïve neonatal mice. Immunohistological staining for macrophages (F4/80; green) and mast cells (MCPT6; red) are indicated. The nuclei are stained with DAPI (blue). **C.** Muscle inflammation with 1% carrageenan increases the number of macrophages and mast cells found in the muscles of neonates at 1d but this is blocked in mice treated with GH. Statistics: F4/80 *p = 0.0006 and ^p = 0.0028; MCPT6 *p = 0.0377 and ^p = 0.0115. One way ANOVA with Tukey’s post hoc multiple comparisons test, n = 4-6/group. Mean ± SEM.

Therefore, we investigated the expression of macrophage and mast cells markers (F4/80 and MCPT6) by immunohistochemistry in the inflamed muscle tissue from our groups. An increase in the numbers of macrophages and mast cells were observed in muscles from inflamed mice at day 1 compared with naïve mice (Figure 3B). The effect was reversed after GH intervention. (Figure 3C). These results suggest that although deletion of GHr from macrophages does not alter inflammatory pain-like behaviors in neonates, muscle inflammation does recruit immune cells into the injury site which can be blocked by GH treatment.

### Nerve targeted knockdown of *STAT1* alleviates pain-related behaviors and macrophage infiltration after muscle inflammation

Results suggested that under conditions of neonatal muscle inflammation, sensory neurons may be playing a significant role in hypersensitivity and possibly inflammation itself in relation to GH-related anti-nociception. We therefore tested a series of TFs that are known to be downstream targets of GH signaling [30] in the DRGs of neonates from our conditions. We found that muscle inflammatory injury markedly increased *STAT1* mRNA expression in the DRGs but not the level of *STAT3, STAT5a, STAT5b* or other GH related TFs, 24 hours post muscle-injection of 1% carrageenan compared to naïve (Table 1). This data is consistent with the gene expression results of mice with cutaneous inflammation (Table S1). Data thus suggests that select TFs may be induced by muscle inflammation in neonates to modulate sensory neuron function.

**Table 1:** Transcriptional changes in L3/4/5 DRGs after muscle inflammation in neonatal mice. STAT1 was significantly upregulated after neonatal muscle inflammation when compared with naive. Data are shown as percent change vs. naïve group with *p < 0.05 vs. Naïve. One-way ANOVA with Tukey’s post hoc multiple comparisons test, n = 4-6/group.

| <u>% Change in DRG mRNA</u> |  |
| --- | --- |
| <u>Gene</u> | <u>1% Carr</u> |
| ELK1 | -23 +/- 9% |
| ELK3 | -4 +/- 12% |
| ELK4 | -19 +/- 12% |
| **STAT1** | **291 +/- 25%\*** |
| STAT3 | 12 +/- 12% |
| STAT5a | 5 +/- 6% |
| STAT5b | 9 +/- 7% |
| SRF | -31 +/- 9% |

We therefore tested whether GH was able to modulate STAT1 expression and found that GH indeed could block inflammation induced STAT1 upregulation in the DRGs (Figure 4A). We then investigated how neuronal STAT1 upregulation may modulate pain-related behaviors after neonatal muscle inflammation. Using our nerve targeted siRNA-mediated knockdown strategy, we first confirmed by RT-qPCR (Figure 4B) and western blotting (Figure 4C) that this injecting Penetratin-1 modified siRNAs against STAT1 into the sciatic nerve was effective at reducing STAT1 upregulation in the DRGs of mice with carrageenan induced inflammation. Consistent with the previous behavioral data, muscle inflammation induced significant paw guarding, mechanical hyperalgesia, and increased paw volume in control siRNA injected neonates (Figures 4D-F). There were no observable changes in the guarding score and paw volume in the contralateral hind paw, but there was a significant restoration of the reduced withdraw threshold observed after muscle injury (Figure S2). Similar to GH treatment, neuronal *STAT1* knockdown, however, blocked nociceptive behaviors observed after muscle inflammation (Figures 4D-F).

**Figure 4:**
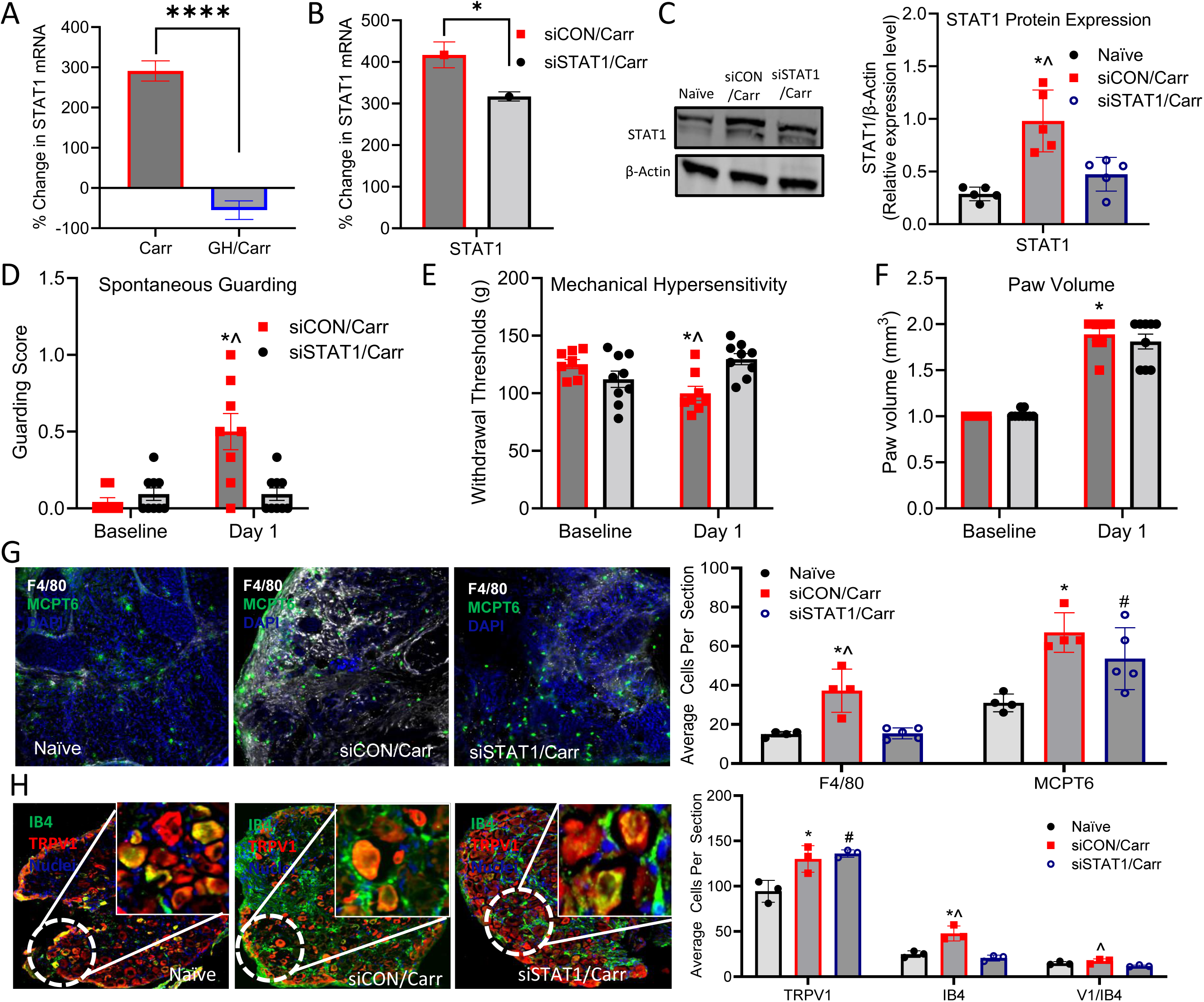
Neuronal STAT1 knockdown alters nociceptive behaviors and prevents infiltration of specific immune cells after muscle injury but has minimal effect on other neuronal markers. **A.** Local GH inhibited the expression of neuronal STAT1 after muscle inflammation in neonates at 1 day. **B.** Confirmation of nerve targeted siRNA-mediated knockdown of STAT1 by real-time RT-qPCR. Data shows percent change from naïve mice. There was a significant increase of STAT1 in the DRGs after neonatal muscle inflammation in control siRNA injected mice (siCON/Carr) and this is significantly reduced by nerve targeted knockdown of STAT1 in mice with muscle inflammation (siSTAT1/Carr) (*p<0.05 vs. siCON/Carr). **C.** Validation of STAT1 knockdown by western blot. Quantification of western blot analysis indicates a significant increase of STAT1 protein expression in the DRGs after muscle inflammatory injury (siCON/Carr) compared to the naïve group, but nerve-targeted siRNA knockdown of STAT1 significantly reduced this expression (siSTAT1/Carr). **D.** Paw guarding scores one day after neonatal muscle inflammation were significantly increased compared with their baseline (BL) but this is not observed in mice with neuronal STAT1 knockdown plus muscle inflammation (siSTAT1/Carr). **E.** Muscle withdrawal thresholds in the same animals showed a significant reduction after carrageenan induced inflammation at 1d compared with BL. siSTAT1/Carr groups showed withdrawal thresholds similar to their baseline at 1d. **F.** There was a significant increase in paw volume in the siCON/Carr group compared to BL, but this effect was not significant in siSTAT1/Carr group. **G.** Neuronal siRNA knockdown of STAT1 selectively inhibits macrophage infiltration (F4/80, white) that is induced by muscle inflammation in neonates at 1d when compared with the naïve group. The inflammation induced increase in mast cells (MCPT6, green) in the injured muscles of neonates at 1d was not inhibited in the siSTAT1/Carr group. The nuclei of the cells were stained with DAPI (Blue). **H.** siCON/Carr mice show a significant increase in TRPV1 (*p=0.0188 vs. naive) and IB4 (*p=0.0054 vs. naive) expression when compared to naive. Also, the inhibition of STAT1 significantly alters IB4 (p=0.6222 vs naïve; ^p=0.0054 vs. siCON/Carr) with no effect on TRPVI expression in the DRG when compared to muscle inflammation in neonates (#p=0.0094 vs. naïve; p=0.6222 vs siCON/Carr). For TRPV1 and IB4 coexpression (VI/IB4), siCON/Carr group showed an increase in VI/IB4 positive cells compared to naïve and siSTAT1/Carr (p=0.3451 vs. naïve; ^p=0.0319), but the reduction observed in siSTAT1/Carr group is not significant when compared to naïve (p=0.2146 vs. naïve). Statistics: for PCR data, percent change vs. naïve group with ****p<0.0001 vs. Carr and *p<0.05, Unpaired t-test. Results are expressed as mean ± SEM, n = 4-6/ group. Western blot data, *p<0.0003 vs. naïve; ^p<0.0040 vs. siSTAT1/Carr but not naïve. 1-way ANOVA, Tukey’s. n=4/group. For behavioral study, two-way RM ANOVA followed by Tukey’s post hoc multiple comparison test. *For comparison to BL; p-values: * ≤0.05. ^for comparison between groups; p-values: ^ ≤0.05 and n = 8-12/ group. For muscle IHC data, F4/80 *p = 0.0148 and ^p = 0.0048; MCPT6 *p = 0.0333 and siCON/Carr vs siSTAT1/Carr, p value = 0.32915. ANOVA with Tukey’s. For neuronal ICH, *p<0.05 vs. naïve, ^p<0.05 vs. siSTAT1/Carr but not naïve and #p<0.05 vs siSTAT1/Carr but not siCON/Carr. 1-way ANOVA, Tukey’s. n=3/group. Mean ± SEM.

Since our GH intervention prevented immune cell infiltration after muscle injury and can modulate STAT1 expression in the DRGs, it was reasonable to hypothesize that neuronal STAT1 may be modulating tissue immune function in part to regulate pain-like behaviors after muscle inflammation. We therefore measured the immune markers F4/80 and MCPT6 in mice with neuronal STAT1 knockdown after muscle inflammation and compared the staining pattern with inflamed mice without STAT1 knockdown (siCON or naïve) (Figure 4G). We detected a significant increase in the F4/80 and MCPT6 positive cells after muscle inflammation in control siRNA injected mice (siCON/Carr) compared to naive. However, in muscle inflamed mice with sensory neuron STAT1 knockdown, the observed increase in F4/80 positive cells was completely blocked. No effect on MCPT6 positive cells in STAT1 knockdown mice was observed, however.

In addition, we quantified the labeling patterns of transient receptor potential vanilloid type 1 (TRPV1-red) and isolectin B4 (IB4-green) after nerve targeted STAT1 knockdown in the DRG neurons to investigate the role of STAT1 in neurochemical alterations after muscle inflammation in neonates [31]. The representative immunohistological staining indicated that there was a significant increase in TRPV1-positive cells after muscle injury compared to naïve. However, STAT1 knockdown was not found to statistically alter the increased expression of TRPV1. Furthermore, there was a small but significant increase in IB4 positive cells after muscle inflammation but this was not observed in the GH treated mice with inflammation. A small number of cells containing both markers (V1/IB4) were found in the DRG neurons of all conditions. Although, the coexpression level of V1/IB4 in siCON/Carr was increased slightly, this was not found to be statistically significant relative to naïve but was different from the STAT1/Carr groups with muscle inflammation. This data suggests that GH may block pain-like behaviors in neonates at least in part by suppressing a STAT1 mediated regulation of macrophage infiltration to the injured muscles.

### GH and *STAT1* inhibition alter neuroimmune signaling in neonatal DRGs after muscle inflammation

Since macrophages appeared to be modulated by both local GH and neuronal STAT1 upregulation, we performed a targeted screen of select factors known to modulate macrophage infiltration and function during injuries in the DRGs from our various groups. We found that the expression of the cytokines monocyte chemoattractant protein 1 (MCP1) and C-X-C chemokine ligand 10 (CXCL10) were upregulated in the DRGs after muscle inflammation while interleukin 1β (IL1β) and CC motif chemokine ligand 27a (CCL27a) were not altered. Interestingly, the upregulation of MCP1 and CXCL10 was blocked by local GH treatment. STAT1 knockdown in neurons also inhibited the upregulation of MCP1 specifically (Figure 5A).

**Figure 5:**
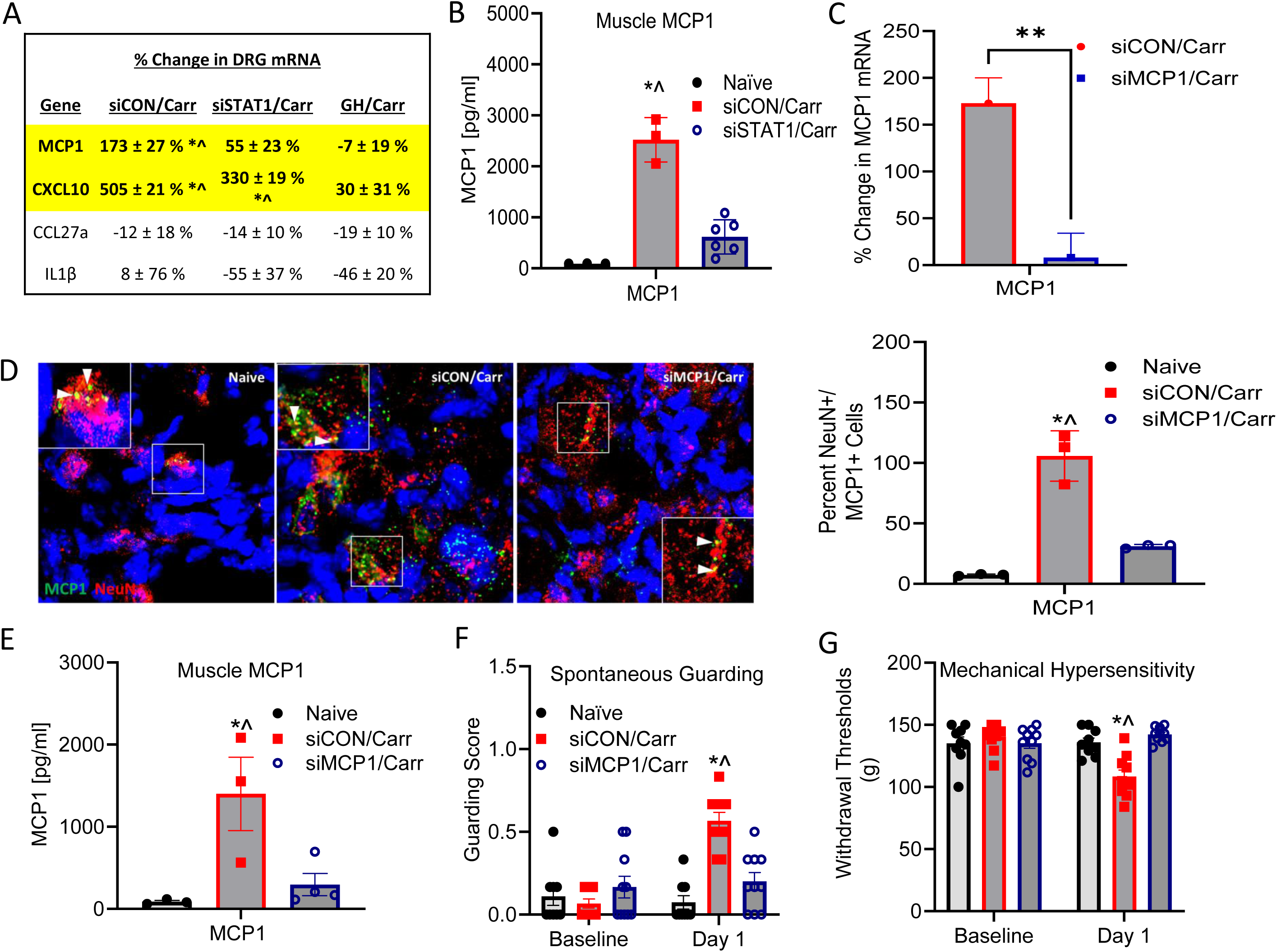
STAT1 regulates MCP1 expression and release from sensory neurons to modulate pain-like behaviors after muscle inflammation in neonates. **A.** Induced neuronal upregulation of select cytokines after muscle inflammation is independently reversed by GH intervention or STAT1 inhibition (siSTAT1/Carr) vs. controls (siCON/Carr). **B.** Muscle MCP1 protein levels as revealed by ELISA were significantly increased 1d after 1% carrageenan injection into the muscles of P7 mice when compared to the naïve group but sensory neuron STAT1 inhibition reversed this effect. **C.** Confirmation of siRNA-nerve targeted MCP1 knockdown by real-time RT-qPCR. Data shows percent change from naïve mice. There was a significant difference between muscle inflammation before and after MCP1 knockdown (**p<0.0067 vs. siCON/Carr). **D.** Validation of MCP1 knockdown by RNAScope. Quantification of RNAScope analysis indicates a significant increase of MCP1 transcript in neurons (NeuN) after muscle inflammatory injury (siCON+Carr) compared to the naïve group (*p= 0.0037 vs naïve) but nerve targeted MCP1 knockdown using siRNAs significantly reduced this effect (^p=0.0179 vs siMCP1/Carr). **E.** MCP1 expression in the muscle was highly upregulated after muscle inflammation in neonates at 1d compared to the naïve group (*p=0.0010 vs naïve) but nerve targeted MCP1 knockdown in inflamed mice (siMCP1/Carr) showed the opposite effect (^p=0.0018 vs siMCP1/Carr). **F.** Nerve targeted MCP1 knockdown in sensory neurons alleviated pain-like behaviors in neonatal mice. Neonatal mice with muscle inflammation displayed increased guarding scores vs BL in control mice (siCON/Carr; *p<0.0001 vs naïve) but not in muscle inflamed mice with neuronal MCP1 inhibition (^p<0.0001 vs siMCP1/Carr). **G.** Muscle withdrawal thresholds in the same animals were significantly reduced compared with naives and the siMCP1/Carr group (*p=0.0003 vs naïve; ^p= 0.0002 vs siMCP1/Carr) and at day one (*p=0.0002 vs naïve; ^p= 0.0001 vs siMCP1/Carr). Statistics: PCR data, *p<0.05 vs. naïve; ^p<0.05 vs. GH/Carr group. One-way ANOVA, Tukey’s. n=5-6/group. ELISA data, *p<0.0001 vs. naïve; ^p<0.0001 vs. siSTAT1+1%Carr but not naïve. 1-way ANOVA, Tukey’s. n=3-6/group. MCP1, monocyte chemoattractant protein. MCP1 gene expression data, Unpaired t test, **p values = 0.0067; Values are n=3-6/group. MCP1 muscle ELISA data: *p=0.0010 vs. naïve; ^p=0.0018 vs. siMCP1+1%Carr but not naïve. 1-way ANOVA with Tukey comparison test. n=3-4/group. Behavioral two-way RM ANOVA followed by Tukey’s post hoc multiple comparison test. *For comparison to BL; p-values: * ≤0.05. ^for comparison between groups; p-values: ^ ≤0.05 and n = 8-12/ group. Mean ± SEM.

As both muscle GH and neuronal STAT1 appeared to regulate MCP1 in the DRGs, we chose to focus on this factor further. With MCP1 playing a role in macrophage mobilization [30], we posited that the upregulation of MCP1 in the DRGs, downstream of STAT1, was contributing to MCP1 protein levels in the inflamed muscles. We confirmed that muscle inflammation increased the levels of MCP1 in the injured muscles of neonates but also found that inhibition of STAT1 in DRG neurons suppressed muscle MCP1 protein levels upregulated in the muscles after inflammation (Figure 5B). Using our nerve targeted siRNA knockdown strategy, we then confirmed that neuronal MCP1 could be successfully knocked down in sensory neurons after muscle inflammation (Figures 5C, D). This corresponded to reduced MCP1 protein expression in the muscle after inflammatory injury (Figure 5E) suggesting that the changes in MCP1 levels in the injured neonatal muscles are significantly modulated by sensory neurons. At the behavioral level, neuronal MCP1 knockdown in inflamed mice also blocked paw guarding scores and inhibited muscle mechanical hypersensitivity (Figure 5F and G).

### MCP1 inhibition alters cytokine activity and regulates macrophage accumulation after neonatal muscle injury

To determine whether neuronal MCP1 modulated inflammation itself in neonates, we performed an unbiased screen of various cytokines and chemokines using protein arrays in mice with muscle inflammation and knockdown of MCP1 (Figures 6A and S5). There were eight significantly upregulated proteins in the inflamed muscles including CD54, IL-1rα, CXCL10, CXCL9, TIMP1, IL-16, CXCL11, and TREM1 that were not significantly affected by sensory neuron MCP1 inhibition when compared with naïve. However, the muscle expression levels of MCP1, M-CSF, CCL12, and CCL5 after muscle injury were significantly inhibited following neuronal MCP1 inhibition (Figure 6B). Equally, MCP1 inhibition in sensory neurons reversed macrophage accumulation following muscle injury and normalized the expression to naive, as indicated by immunofluorescence (Figure 6C). No effects of MCP1 inhibition were observed on the increase in mast cells found in the inflamed muscles of neonates. These data suggest that MCP1 is released from sensory neurons in neonates with muscle inflammation to regulate macrophage infiltration and pro-inflammatory signaling in the muscles which contributes to pain-related behaviors.

**Figure 6:**
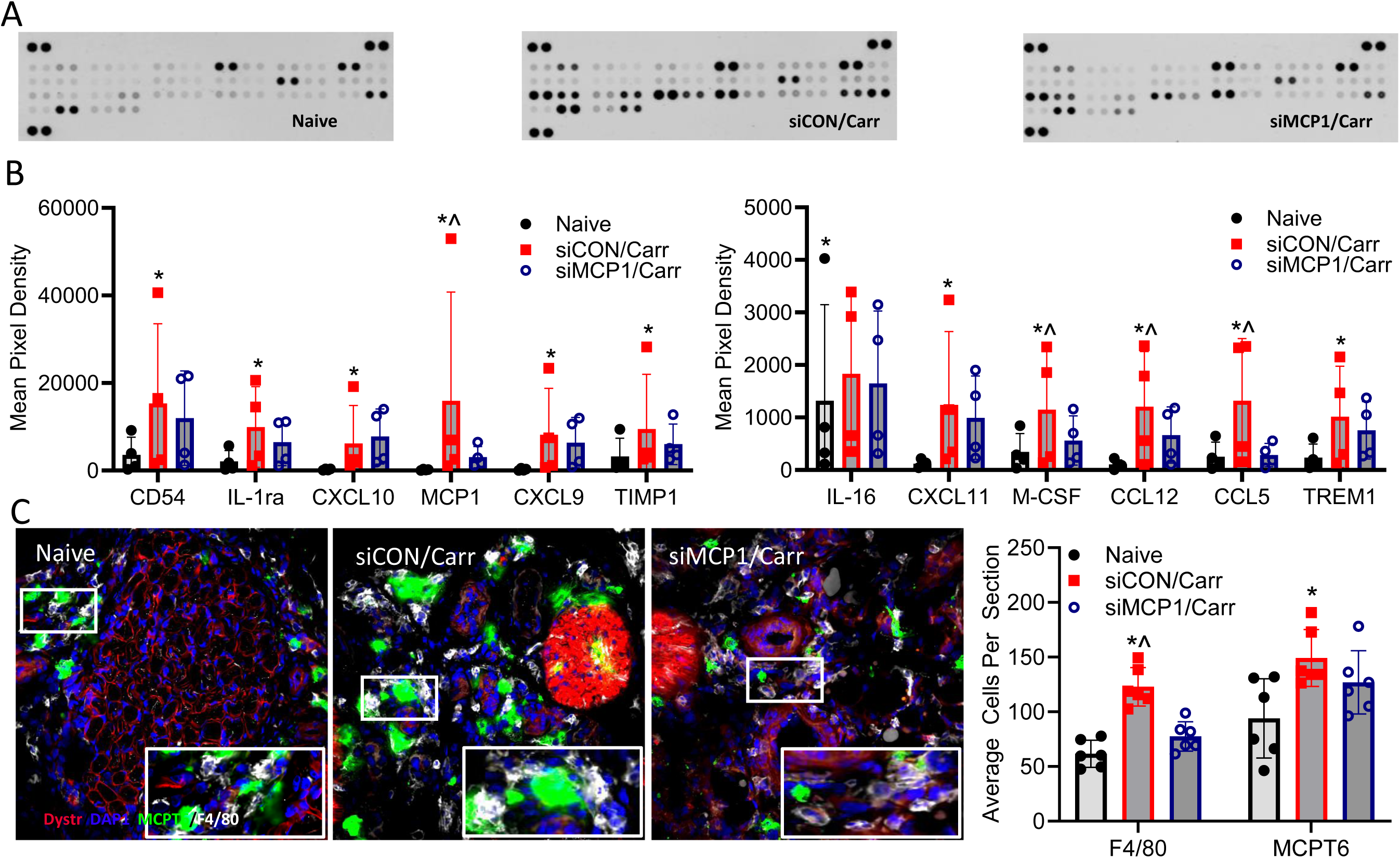
Neuronal MCP1 regulates immune function in the muscles after inflammation. **A.** Representative cytokine/chemokine protein arrays processed from muscles of naive, siCON/Carr, and siMCP1/Carr conditions. **B.** Quantification of the mean pixel density in differentially expressed proteins under the different conditions indicated. Proteins with significant differences from the naïve condition were represented. The expression of MCP1, M-CSF, CCL12, and CCL5 after muscle injury were significantly downregulated by MCP1 inhibition in neonates with muscle inflammatory injury. **C.** Neuronal MCP1 inhibition inhibits macrophage (F4/80, white) infiltration into the muscles after inflammation at 1d but this does not alter changes in mast cells (MCPT6, green) observed in the muscles of inflamed mice. DAPI (blue) is used to mark nuclei and dystrophin (red) indicates muscle fibers. Statistics: For cytokine array, *p < 0.01 vs Naïve and ^p < 0.01 vs siMCP1/Carr. Two-way ANOVA Turkey’s (Mean ± SEM; N = 3-4/ group). For muscle IHC data, *p=0.0016 vs Naïve and ^p <0.0073 vs siMCP1/Carr. One way ANOVA Turkey’s comparison test; N = 3/group.

## Discussion

The body responds to tissue injury with inflammation to initiate the healing process. This is orchestrated by a cascade of events involving immune cells and inflammatory mediators, which also triggers inflammatory pain by sensitizing and activating sensory nerve fibers [32]. Inflammation increases sensitivity to various stimuli, making it challenging to treat therapeutically. GH and its related signaling molecules play an important role in neonatal nociception and development of chronic pain [33]. GH dysregulation has been linked to hyperalgesia in common clinical pain syndromes including fibromyalgia [34] and erythromelalgia [35]. Pretreatment of GH has been reported to reverse mechanical and thermal hypersensitivity in an animal model of neonatal cutaneous inflammatory pain [36], but the mechanism by which this may occur after muscle inflammation was not fully known. In this study, we found a novel mechanism by which muscle GH modulates STAT1 expression in sensory neurons to regulate peripheral sensitization following muscle inflammatory injury. This was found to modulate MCP1 release from sensory neurons which in turn regulated macrophage infiltration to the inflamed muscles and subsequent behavioral hypersensitivity.

Inflammation or injury related aberrant sensory input during crucial stages of early postnatal development can have long-term negative effects on pain perception [37]. GH signaling during early life has been shown to modulate acute pain-like responses after incision and prolonged hypersensitivity after repeated incision [27]. Here we also found that GH intervention at the time of muscle inflammation alters nociceptive behaviors (Figure 1) and inhibits sensory neuron sensitization (Figure 2). In the current report however, we found that unlike after neonatal incision, muscle inflammation selectively induces STAT1 upregulation in the DRGs of neonates which can be blocked by local GH treatment (Table 1). This interestingly was not due to significant alterations in expression of channels such as TRPV1 (Figure 4) but was due to its modulation of cytokine release (MCP1) from sensory neurons (Figures 3, 5 and 6). Evidence suggests that GH signaling may be an important global regulator of somatosensory development in neonates [8]. This is supported by clinical evidence that GH may regulate pain in patients with GH deficiency [35]. Our new evidence, however, suggests that GH can uniquely engage distinct sensory neuron related mechanisms of pain development in neonates depending on the injury sustained. This will need to be critically evaluated in future studies that assess distinct injuries across the lifespan.

Calcium transients critically regulate numerous neuronal functions, including excitability, kinase activity, neurotransmitter release, gene expression, and apoptotic cell death [38]. Here we utilized a new neonatal *ex vivo* muscle/nerve/DRG/spinal cord Ca^2+^ imaging preparation to assess neuronal activity in the DRGs using GCaMP6 after muscle inflammation. (Figure 2). This preparation is advantageous as it allows for the acquisition of functional activity in several cells simultaneously in response to muscle RF stimulation with various sensory modalities. We found that the force required to induce calcium responses in DRG neurons in mice with inflammatory injury was lower compared with naive and inflamed mice treated with GH. In addition, heat sensitive cells were increased in mice with muscle inflammation compared to naïve. This effect was reduced by GH intervention. Although there were no significant changes in the number of mechanically sensitive cells, those that responded to heat stimulation of the RFs and afferents that responded to both metabolite mixtures were found to be significantly higher relative to naïve conditions. This was reversed in mice with muscle inflammation and the GH intervention. This pattern of sensitization is somewhat distinct from that observed using single unit recordings after incision injury [27] in that here we additionally found an alteration in the heat sensitive fibers. This supports the notion that distinct neonatal injuries can produce pain like behaviors through unique mechanisms. It is important to note that we are not able to separate the cells by conduction velocity with this preparation. However qualitative assessments of calcium florescence in our conditions suggests that both myelinated and unmyelinated afferents contribute to the results obtained here (Figure 2). Nevertheless, other reports have shown alterations in afferent heat sensitivity under conditions of inflammation in adults (Jankowski et al 2012), and here we show that GH appears to be able to modulate this afferent sensitization pattern in neonates as well but not necessarily by regulating the numbers of cells expressing TRPV1 (Fig. 4).

### Targeting STAT1 expression in DRG neurons suppresses nociceptive behavior and immune cell infiltration

GH is known to activate a variety of TFs in cells such as ERK-like kinases (ELKs), STATs [11], and SRF [39] to name a few [40]. Of all of these however, only STAT1 was overexpressed in the DRGs after muscle inflammation at 24 hours in P7 neonatal mice (Table 1). This expression was directly involved with increased pain-like behaviors (Figure 4). Walker et al., showed that STAT1 overexpression is correlated with inflammatory arthritis in synovial tissue biopsy specimens from patients with rheumatoid arthritis, spondyloarthritis, and osteoarthritis compared to normal tissue [41]. Since STAT1 is a major mediator of cellular responses and plays a crucial role in immune cell function, we investigated the involvement of neuronal STAT1 on immune infiltration. Macrophage and mast cell infiltration after muscle inflammatory injury result in an immune cell-dependent GH regulation (Fig 3 C and D). Although other TFs not analyzed here could play a role in neonatal inflammatory myalgia, local GH was found to at least suppress neuronal STAT1 expression which regulated immune cell mobilization after muscle inflammation and consequently modulated peripheral sensitization.

STAT1 knockdown in the sensory neuron after inflammatory injury prevented behavioral hypersensitivity and immune cell infiltration in the muscle with minimal effects on TRPV1 and IB4 in the DRGs (Figure 4). One study reported that the inflammatory impact of IFN-γ in CD8+ T is partly mediated by STAT1 pathway in lung injury induced by LPS [42]. The immunohistological changes in the muscles observed after STAT1 knockdown correlate well with the effect shown by GH intervention (Figures 3-4). Immune cells modulate nociceptors to alter pain sensitivity [43]. Studies show that macrophages and neutrophils initiate the release of inflammatory cytokines to promote repair [44]. Our result showed that the elevated immune cells caused by muscle inflammation were reversed independently by local GH delivery or neuronal STAT1 knockdown (Figures 1 and 4). Thus, under particular injury conditions in neonates, TFs upregulated in sensory neurons may modulate hypersensitivity through their control of inflammation itself in the injured target. Together, STAT1 knockdown may be sufficient to decrease inflammatory pain and targeting STAT1 expression may be a novel approach to reduce pain in neonates.

### GH and STAT1 inhibition regulate neuroimmune signaling in mice with muscle inflammation

Reports have shown acute inflammation induced STAT1-dependent chemokine secretion in tubular epithelial cells [45]. Our study showed that the expression of neuronal MCP1 and CXCL10 were higher after muscle injury, an effect that was independently suppressed by either GH or STAT1 knockdown (Fig. 5A); STAT1 having a specific effect on MCP1 levels. To determine if STAT1 regulated MCP1 secretion into the muscles, we measured protein level of MCP1 in muscle supernatant after injury and its corresponding effect after STAT1 knockdown (Fig. 5B). We found that virtually all of the changes in MCP1 protein induced in the muscles by inflammation were due to changes in STAT1 and MCP1 upregulation in neurons (Figures 5). Although MCP1 has been shown by countless studies to be involved in immune cell recruitment and inflammation [46–48], here we show a novel role for neonatal primary afferents in the production of this cytokine during muscle inflammation.

This is further supported by data showing that both STAT1 inhibition and MCP1 knockdown in neurons of neonates with muscle inflammation blocks the infiltration of macrophages to the injured tissue (Figures 4-6). This in turn modulated the inflammatory cytokine profile detected in the neonatal muscles (Figure 6). Certain chemokines are highly expressed under certain conditions including inflammation [49], and peripheral nerve injury [50]. Their receptors are expressed on the somatosensory neurons [51] which can regulate hypersensitivity [52]. Data shows that MCP1 receptor knockout (CCR2) prevents hypersensitivity in mice with the partial sciatic injury [53]. Inflammatory cytokines orchestrate peripheral inflammatory responses to alter behavioral activities including increased pain [54]. Inhibition of MCP1 in the DRG prevented behavioral hypersensitivity and proinflammatory signaling in the muscles (Figures 5-6). This could indicate that targeting neuroimmune signaling is a novel strategy for treating pain and inflammation in neonates.

Recent reports have begun to show that sensory neurons are key drivers of inflammation [55–57]. MCP1 has been a recent cytokine found to be expressed in sensory neurons that may be involved in pain responses in adult injury models [58, 59]. Our data now shows that neonates may utilize a unique non-peptidergic, neurogenic inflammation to modulate hypersensitivity after muscle inflammation. Results could uncover new ways to treat pain or inflammation itself in children with distinct types of muscle damage.

## Author contributions

A.O.F., A.J.D., and M.P.J conceptualized the research study; A.O.F., and M.P.J developed the methods to be used A.O.F., M.P.J, P.G., N.G.R.R, and M.C.H. performed the experiments outlined; A.O.F., and M.P.J analyzed the data; A.O.F. wrote the initial draft of the manuscript with editing and further writing by M.P.J.

## Acknowledgements

This work was supported by grants from the NIH (R01NS105715 and R01NS113965) to MPJ and a fellowship to AOF from the Cincinnati Children’s Research Foundation (FP00013896). We would like to thank Dr. Xinzhong Dong for generously donating the Pirt-Cre mice used in these studies to generate the sensory neuron calcium reporter line (PirtCre;GCaMP6f).

## Methods

### Animals

Postnatal day 7 (P7) male and female Swiss Webster, C57BL/6J, PirtCre;GCaMP6f or LysMCre;GHr^f/f^ mice along with their littermate controls were used in these studies. All animals were housed in the CCHMC barrier facility and maintained under standard laboratory conditions (14 h light/ 10 h dark cycle, lights on at 06:00h, and a temperature-controlled environment) and food and water available *ad libitum* for the parents. All procedures were approved by the Cincinnati Children’s Hospital Research Foundation Institutional Animal Care and Use Committee and adhered to National Institutes of Health Standards of Animal Care and Use under practices approved by the Association for Assessment and Accreditation of Laboratory Animal Care. Animals were anesthetized with either 2-3% isoflurane before induction of various treatments or with ketamine/xylazine (100 mg/kg / 10 mg/kg) prior to dissection for terminal procedures.

### Carrageenan-induced inflammation

Under isoflurane anesthesia, 1% Carrageenan (in 0.9% NaCl) was injected into the right hairy hind paw skin or right hind paw muscles of neonatal mice and compared to needle puncture with no injection of substance (sham) or naive controls. For interventions, co-injection of 10 µL of 1%Carr+1.5 mg/kg mouse recombinant growth hormone (GH) or vehicle (Carr+saline) was injected into the right hind paw muscle 1d after baseline behavioral studies. Prior to and post inflammation, the body weight and the hind paw volume measurements were recorded using the weighing balance in grams and Vernier calipers (mm^3^), respectively.

### Pain-related Behavioral Assessments

Behavioral examination of spontaneous paw guarding, and mechanical withdrawal thresholds to muscle squeezing were performed in the groups described. Before the commencement of the behavioral studies, mice were randomly assigned to different groups using a random number generator into a rectangular transparent plastic box after which they were acclimatized for not more than 10 minutes to avoid keeping the pups away from their parents for a significant period of time. Baseline pain scores were measured before the induction of inflammation. Behavioral assays were conducted by experimenters blind to conditions.

#### Spontaneous paw guarding

Pups were allowed to move freely inside the transparent plastic box. The ipsilateral and the contralateral paws were closely monitored during a 1 min period repeated every 5 min for 30 minutes. The testing scale scores paw movement ability ranges from where there is no observable hind paw movement (score = 0), through when a slight raise is observed (score = 1) to when the paw is completely raised (score =2). Guarding was evaluated one animal at a time at 5-minute interval for six readings and the average score is used as the mean value.

#### Mechanical withdrawal thresholds

A digital paw pressure device with a modified, rounded blunt tip was used to quantify the withdrawal thresholds to muscle squeezing before and 1d after muscle inflammation with or without various interventions described. This device was used to apply increasing mechanical force (up to 150 grams) on the plantar surface of both hind paws until a paw withdrawal was observed. This was repeated three times with at least a 5-minute interval between trials. The average of the trials was recorded per mouse and the average per condition was calculated and compared as noted.

### Gene expression: RNA extraction and quantification by real-time RT-PCR

Messenger RNA (mRNA) was extracted from L2-3 (for cutaneous inflammation conditions) or L3-5 DRGs (for muscle inflammation conditions) from P7-P8 pups using RNeasy Mini Kits (Qiagen Stock#: 74104) and the mRNA was DNase I treated, and reverse transcribed into cDNA using the SuperScript™ II Reverse Transcriptase for RT-PCR (cat. no. 18064022, Invitrogen, Carlsbad, CA, USA) according to the procedure provided by the manufacturer. The expression levels of the transcription factors, cytokines, and other genes indicated were quantified by SYBR Green Master Mix using real-time quantitative RT-PCR with an Applied Biosystems™ StepOnePlus™ Real-Time PCR machine (Applied Biosystems™ 4376598, Fisher scientific). The primer sequences for the targeted genes in this study were designed and tested for efficiency using Primer-BLAST tool in NCBI at http://www.ncbi.nlm.nih.gov/tools/primer-blast/ (Table S2). The assays were performed in two replicates, and the results were normalized to the internal standard mRNA levels (GAPDH) using the formula 2^ΔΔCT^. Data were analyzed on the raw normalized PCR values, but data were presented as percent change in expression relative to control.

### siRNA knockdown experiments

#### Analysis of knockdown efficiency in vitro

To determine the siRNA duplexes with the highest knockdown efficiency to be used in the main experiments of this study, we first determined this in mouse neuroblastoma cell-lines (Neuro2A cells: CCL-131, Lot No. 60279356) purchased from American Type Cell Culture (ATCC, USA). Neuro 2a cells were transfected with non-targeting siRNA or one of 4 different siRNAs targeting either STAT1 or MCP1 individually (Dharmacon™ siRNA solutions). Neuro2A cells were cultured in complete growth medium (Eagle’s Minimum Essential Medium, Catalog No. 30-2003) supplemented with 10% Fetal bovine serum and 1% Penicillin-Streptomycin Gibco™ (Catalog No. 15070063) for 24 hrs (37 °C in a 5% CO_2_ and 95% humidified atmosphere). Prior to siRNA treatment, cells were subcultured in 12 well culture plates at a density of 1×10^4^ cells/ml and allowed to reach 70% confluency. Cells were transfected using 10 nM of siRNAs with 2.5 µl TransIT-TKO® Transfection reagent (Mirus) for 24 hrs following manufacturer’s recommended protocols. The knockdown efficiency of the siRNAs was determined by real-time RT-PCR after RNA isolation using RNeasy Mini Kits (Qiagen) and cDNA synthesis by SuperScript II. The mRNA levels of each of the duplexes of STAT1 and MCP1 tested in cell cultures were compared with the mRNA levels of the non-targeting siRNA-treated conditions. The duplex with the highest knockdown efficiency was used for further in vivo analyses.

### Modification and conjugation of siRNAs

STAT1 and MCP1 siRNAs with the sense and antisense sequences 5’-GCAUAGAGCAGGAAAUCAA-3’ and 5’-GAAGUUGACCCGUAAAUCU-3’, respectively, were modified using the ‘Designed modified siRNA’ tool available within Dharmacon homepage (https://horizondiscovery.com/en/gene-modulation/knockdown/sirna). The sense and the antisense strands were 5’ thiolated and 5’ phosphorylated, respectively, with 3’-UU overhangs. The modified siRNAs were thiol deprotected using TCEP as a reducing agent according to the manufacturers protocol and subsequently conjugated with lipophilic peptide Penetratin-1 (Pen-1: MP Biomedicals). Briefly, the modified annealed duplexes were dissolved in 400 μL 3% Tris (2-carboxyethyl) phosphine followed by the addition of 50 µl of 3 M sodium acetate and 1.5 mL of 200 proof ethyl alcohol. The resulting mixtures were incubated at –80 °C for 20 minutes and centrifuged at 13000 × g for 20 minutes (4 °C). To the pellets, 200 μL of 95% ethanol was added and immediately discarded and further dried in speed-vacuum centrifuge. The dry siRNA pellets were then resuspended in 1x siRNA buffer. Furthermore, 25 μl of 2 mg/mL Pen-1 was added to each tube and the mixtures were heated to 65°C for 15min and then incubated at 37°C for 1 hour. 5M NaCl was then added to the final solution and stored at -80°C prior to nerve injection. This process was also carried out with non-targeting siRNAs (Thermo Fisher Scientific, catalog #D-001206-14-05), indicated as siCON.

### Nerve-specific siRNA injections

The sciatic nerve was carefully exposed in the right hind limb above the knee, proximal to the trifurcation before being lifted onto a Parafilm platform for siRNA injections at P6, 1d prior to muscle inflammation or baseline behaviors. The sciatic nerve was then pressure-injected at 1-2 psi with ∼ 0.2 µl of prewarmed 90 µM Pen-1 conjugated siRNAs (siSTAT1, siMCP1 or siCON) using a quartz microelectrode and picospritzer. The injection site was closed using 7.0 silk sutures after siRNA injection and parafilm removal.

### Immunocytochemistry and volume imaging

DRGs and muscles were processed either by cryostat sectioning or whole tissue imaging. Prior to these processes, P7 neonatal mice were briefly anesthetized with 2-3% isoflurane followed by intramuscular injection of Ketamine/Xylazine Solution (9 mg/mL Ketamine + 0.9 mg/mL Xylazine) to induce deep anesthesia and perfused with ice-cold saline. DRGs were harvested in either OCT medium on dry ice or directly in ice-cold 4% paraformaldehyde (PFA). Muscle tissues were harvested without perfusion in either OCT medium on dry ice or directly in PFA. Cryomold samples were sectioned (14 µm thick) and prepared on a cryostat. Sectioned samples were washed and blocked with gelatin blocking buffer in 1x PBS for 1 hr, followed by overnight incubation at 4 °C with the following primary antibodies: rat anti-F4/80 (1:500; Abcam ab6640) and goat anti-MCPT6 for muscle tissues or rabbit anti-TRPV1 (1:200; Alomone Cat No. ACC-030) and IB4-647 (1:1000; Invitrogen) for DRG neurons.

For volume imaging, tissues were fixed in 4% PFA overnight at 4 °C and delipidated in 100% methanol for 1 hr on ice. Tissue permeabilization was carried out in dent’s bleach (4:1:1 MeOH: DMSO: 30% H_2_O_2_) for 2 hrs at room temperature followed by treatment with graded methanol solutions (100%, 75%, 50%, 25%, each for 30 minutes at 4 °C). The tissues were then rinsed twice in 0.01 M PBS buffer for 1 hour. After bleaching, tissues were washed in EZ away solution (tetrahydrofuran: water) overnight at room temperature followed by four rinses with milli-Q H_2_O for 1 hour each at 37 °C and then incubated in primary antibodies. Prior to detection, samples were washed and incubated with secondary antibodies including Alexa fluor-647 donkey anti-rat, Alexa fluor-594 donkey anti-rabbit, Alexa fluor-488 donkey anti-goat, Cy3 fluorophores, and 647-Conjugated (1:400; Jackson ImmunoResearch Laboratory). Slide sections were cover-slipped with mounting media prior to imaging while whole tissues were mounted with Refractive index (RI) matching solution on glass slides with rings shims. Nuclei were stained by DAPI solution for 15 minutes at room temperature. Fluorescence images were captured with Nikon AXR Upright Confocal Microscope with either 16x LWD objective or 20/40x Plan Fluor Multi-Immersion Objective. Quantitative analyses of immunofluorescence staining images were carried out by ImageJ and Nikon NIS element imaging software.

### Protein analyses

The DRGs or hindpaw muscle of neonatal mice treated with 1% carrageenan or GH/ Carrageenan or siRNA p6 nerve injection with inflammation were first processed for protein quantification. Briefly, mice were anesthetized and the right hind paws of were dissected without perfusion. After skin removal, paw muscle samples were kept on dry ice if not processed immediately or homogenized at 4°C in protein lysis buffer (50 mM Tris pH 7.4, 0.5% SDS, 1 µg/ml leupetin, 1 µg/ml Aprotinin, 1 µg/ml Pepstatin, 1 mM sodium orthovanadate, and 1 mM PMSF). For DRGs, mice were perfused before DRG dissection prior to homogenization. The homogenates were centrifuged at 12000 rpm for 15 min at 4 °C. The supernatants were transferred into a new collection tube and stored at -80°C for downstream analysis. Protein quantification was determined against bovine serum albumin (BSA) standard with a detergent compatible (DC) protein assay kit (23225, Thermo Fischer Scientific).

### Enzyme-linked immunosorbent assay (ELISA)

Twenty-four hours after 1% carrageenan injection, the mice were anesthetized, and the right-hind paws were dissected and stored in dry ice in labeled tubes and then individually homogenized after the removal of the skin. The levels of MCP1 in the muscle lysate were measured by commercial MCP-1 Mouse ELISA Kit (Catalog No. BMS6005, Invitrogen Thermo Fischer Scientific). Briefly, 200µg protein lysates were diluted with assay diluent provided and processed according to the manufacturers’ instructions. The diluted lysates were incubated on the capture antibody-coated plates with biotin-conjugate for 2 hours at room temperature. The standard sample and blank were processed in the same fashion and all samples were performed in duplicates. Next, all unbound proteins were removed by washing each well three times with wash buffer following the incubation of streptavidin-HRP for 1 hour at room temperature. Prior to detection by spectrophotometer at 450 nm, 3,3’,5,5’ tetramethylbenzidine substrate solution was added to all the wells and incubated for 10 minutes at room temperature in the dark after which the enzymatic reaction was terminated using the stop solution. Unknown muscle protein lysates were extrapolated from the MCP1 standard curve using Graph pad prism v9.6 and the results were expressed in units of pg/ml.

### Western blot

The L3-5 DRGs from 3 pups were pooled as n=1 to obtain a significant protein concentration and three pooled samples per condition were analyzed across conditions. The protein lysate/ supernatants were quantified and protein expression in the muscle after neuronal STAT1 knockdown was determined by western blot. Briefly, 30 µg of protein lysates were added to 5x Laemmli buffer and heated to 95 °C for 10 minutes following the formula: 30 µL – 4 µL SB – Z µL lysate = Y µL H_2_O. 20 µl of the mixtures were resolved along with Chameleon® Duo Pre-stained Protein Ladder (Lot #928-60001) in 10% precast Sodium dodecyl-sulfate polyacrylamide gel electrophoresis (SDS-PAGE: Bio-Rad, cat# 456-1030). Samples were separated at 50 V for 30 minutes followed by 100 V for 1 hour. Protein samples were further electrophoretically transferred from the SDS-PAGE gel to a nitrocellulose membrane (Bio-Rad) at 35 V at 4 °C overnight or at 90 V for 2 hours using the wet electrotransfer method. Membrane washing and antibody incubations were carried out using Invitrogen iBind Automated Western System procedure with iBind kit (Lot #1502853), iBind card (Bi01123, and iBind devise (Lot # 1312006, Novex, life Technologies Corporation, Israel) following the manufacturer’s instruction. The primary antibodies for STAT1 (Cat # 9172L, 1:200 (v/v) dilution), and β-actin (Cat # 3779, 1:500 (v/v) dilution) were incubated along with the secondary antibodies conjugated with IRDye Infrared Fluorescent Dyes (1:1000 (v/v) diluted anti-rabbit for STAT1, and β-actin, with emissions at 800 nm and 680 nm, respectively (LI-COR Biosciences, USA). Detection was achieved by LI-COR Odyssey CLx Infrared Imaging System. Membrane bands were quantified within the LI-COR software using the analysis module. Protein concentrations were further validated using Image J software (Image Processing and Analysis in Java, National Institute of Health) by calculating relative densities of the protein bands.

### Proteome Profiler Mouse Cytokine Array

Muscle samples from the MCP1 knockdown conditions (naïve, siCON/carr, and siMCP1/carr) were dissected out from the right hind paw and kept on dry ice and further stored at −80°C prior to protein extraction and quantification. Cytokines were analyzed using proteome profilers (R&D Systems mouse cytokine array panel A, no. ARY006) loaded with 200 μg of protein samples following manufacturers instruction. Each sample was incubated with a separate array precoated with 40 cytokine/chemokine duplicate antibodies and labeled with IRDye 800CW Streptavidin-conjugated secondary antibody. The intensity of the antibodies was analyzed using Odyssey® CLx Infrared Imaging System by LI-COR. Duplicates were averaged, and the background subtracted to calculate the mean pixel density for each protein.

### Ex vivo calcium imaging

Neuronal calcium activity was studied in mice expressing an endogenous calcium indicator in sensory neurons (PirtCre;GCaMP6f). Both male and female pups were used for calcium imaging studies at postnatal day 7 (three pups per condition). Pups at P6 were injected with 1% carrageenan to induce inflammation or co-injected with GH and 1% carrageenan and at P7. Ca^2+^ transients as a surrogate of neuronal activity were evaluated using an in-house novel ex vivo neonatal hindpaw muscle, tibial nerve, DRG, spinal cord preparation similar to that described for single unit recordings previously [27].

Briefly, pups were dissected in ice cold oxygenated (95% O_2_ / 5% CO_2_) artificial cerebral spinal fluid (aCSF; 127.0 mM NaCl, 1.9 mM KCl, 1.2 mM KH_2_PO_4_, 1.3 mM MgSO_4_, 2.4 mM CaCl_2_, 26.0 mM NaHCO_3_, and 10.0 mM D-glucose: O_2_ aCSF) and the hindpaw muscle, tibial nerve, L1-L6 DRGs, spinal cord were isolated in continuity and then transferred to a recording dish containing oxygenated aCSF (32°C) mounted on the stage of the upright confocal microscope. Imaging was performed separately on L3 and L4 DRGs before (baseline) and after stimulations of the receptive fields. For mechanical stimulation, a series of Von Frey filaments ranging from 0.07 g to 10 g were delivered to 6 distinct RFs in the hindpaw muscles followed by thermal stimulation where ice-cold (∼ 1°C) or hot (∼ 50°C) saline were administered onto the hindpaw muscles. Thereafter, chemical stimulation was tested on the muscle using oxygenated low concentration of metabolites (15 mM lactic acid, 1 µM ATP, pH to 7.0) and then a high concentration of these metabolites (50 mM lactic acid, 5 µM ATP, pH to 6.6) in aCSF. Both mechanical and thermal stimulation were repeated post chemical stimulation.

Neuronal activities were visualized using Nikon AXR Upright Confocal Microscope with 16x LWD objective. Data analysis of imaging recordings of Ca^2+^ transients was performed with Nikon NIS-elements imaging software and ImageJ (National Institute of Health, MA, USA). Recordings were processed to generate maximum florescence intensity over time. Regions of interest (ROIs) were automatically detected in cells that showed an increase in fluorescence intensity. To quantify fluorescence changes in individual cells, values were normalized to baseline and expressed as ΔF/F_O_ with the formula ΔF/F_O_ = (Fmax – F_O_) / F_O_ where Fmax: maximum intensity for stimulation and F_O_: average intensity of the baseline prior to stimulation. The percent cell responders to each of the stimulations (mechanical, heat, cold, chemicals (High, and/or low)) were calculated by dividing cells that responded to stimulation (observable changes in the florescence intensity) by the total number of cells in the DRG multiplied by 100%. The total cell number was determined by creating the maximum intensity projection over time (MaxIP) from all the stimulation videos prior to the detection of the region of interest.

### RNAScope In Situ Hybridization

DRG samples of all conditions (Naïve, Carrageenan only/ siCON/Carr, and siMCP1/Carr) were harvested into chilled 0.1% DEPC PBS and subsequently fixed in 4% PFA at room temperature for 2 hours after which samples were embedded in OCT before storage at -80°C. Frozen tissues were sectioned at a thickness of 14 μM using a cryostat. For sample processing, sectioned slides were dehydrated in an ethanol series and further air-dried for 5 min. Second, following creation of a hydrophobic barrier on the slides, DRG sectioned slides were treated with hydrogen peroxide for 10 mins at room temperature. Third, DRGs were digested in Protease IV for 30 minutes at 40 °C. Lastly, slide sections were treated with RNAscope reagents according to manufacture protocols. Briefly, slides were hybridized in 1:50 (probe to diluent) of MCP1-C2 and Mm-Rbfox3-C3 target probes and incubated in a HybEZ oven (ACD) for 2 hours at 40 °C followed by sequential amplification with AMP1-3 for 30, 30, and 15 mins, respectively at room temperature. Before adding each AMP reagent, samples were washed twice with 1x washing buffer (cat. no. 310091 ACDBio). The HRP-C2 and C3 signals were sequentially developed with Opal Polaris 520 and 620, respectively. The samples were then counterstained with DAPI for 30 seconds at room temperature and cover slips were mounted by ProLong Gold Antifade Mountant prior to imaging on a Nikon A1 inverted confocal microscope.

### Data Analysis

Data were analyzed using GraphPad prism v9.5. Values were normalized and inspected for normal distribution using a Shapiro-Wilk normality test. Results were statistically significant when p < 0.05. All data are reported as meansLJ±LJstandard error of the mean (SEM). Calcium fluorescence was quantified by NIS elements software and analyzed by parametric one-way ANOVA and Tukey post-hoc. All behavioral assays were analyzed via a one or two-way repeated measures analysis of variance (RM-ANOVA) with Tukey’s post hoc tests. For multiple comparisons including muscle and DRGs immunohistochemistry/ volume imaging, western blot, gene expression, ELISA, or cytokine array data, a one or two-way ANOVA with Tukey’s multiple comparisons test was performed according to experimental design. For a two-group comparison such as siRNA knockdown, PCR data as indicated in the figures, an unpaired t-test was performed. Graphical abstract was generated using BioRender.

## Ethical Permissions

All procedures were approved by the Institutional Animal Care and Use Committee at Cincinnati Children’s Hospital Medical Center, under AAALAC approved practices.

## Supplementary Figures

**Figure S1:**
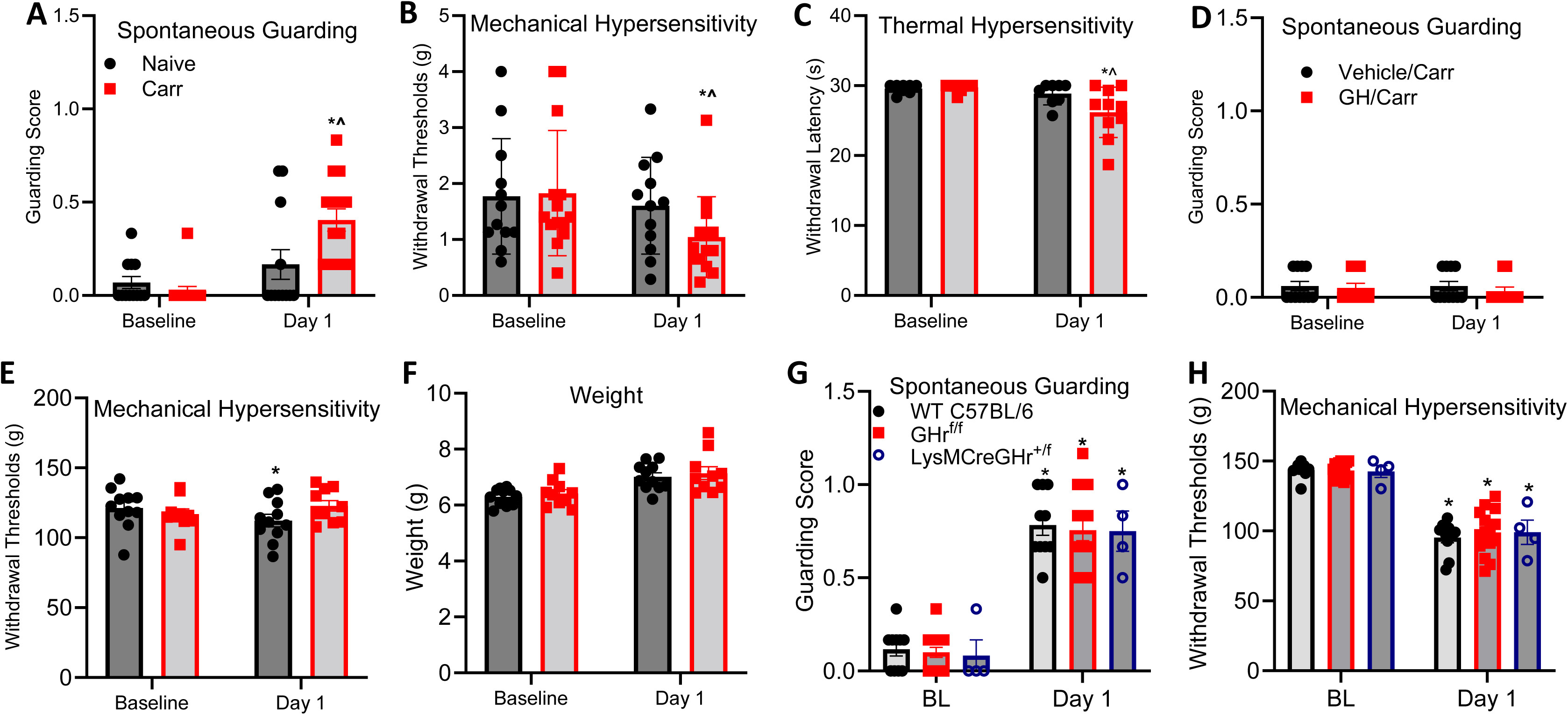
Carrageenan induces cutaneous inflammatory injury and modulates contralateral muscle mechanical hypersensitivity in neonatal mice. **A.** Spontaneous paw guarding revealed a significant difference between carrageenan injected and naïve/sham groups at BL and day 1 but no difference between BL and day 1 in naïve/sham mice (*^p<0.001 vs. naïve at BL and day 1, respectively). **B.** Mechanical hypersensitivity in the muscle inflamed group showed a significant decrease in withdrawal thresholds when compared with naïve/sham and BL while there was no significant difference between naïve/sham group vs BL (*p=0.05 vs. BL). **C.** Mice with cutaneous inflammation revealed lower heat withdrawal latency when compared with BL and naïve at day 1. **D.** Spontaneous paw guarding at BL and Day 1 contralateral to muscle inflammation showed no significant difference between carrageenan injected and naïve/sham groups (*p<0.001 vs. BL for naïve/sham and Carr). **E.** Mechanical hypersensitivity in the contralateral limb post muscle inflammation showed a significant decrease in withdrawal thresholds when compared with naïve/shame and baseline while there is no significant difference between naïve/sham group vs BL (*p=0.05 vs. BL). **F.** There were no changes in weight after carrageenan (Carr) or GH/Carr treatment in neonates at day 1. **G.** No significant differences were observed between groups at BL and/or Day 1 for paw guarding. However, there was a significant increase in spontaneous guarding activity from BL to Day 1 within each group (*p < 0.0001vs. BL). **H.** When assessing mechanical hypersensitivity to paw squeezing, no significant differences were found between groups at BL and/or Day 1. However, all groups exhibited a significant decrease in mechanical withdrawal threshold from BL to Day 1 (*p < 0.05 vs. BL). 2-way ANOVA with Tukey’s post hoc test. n=4-15. Mean ± SEM

**Figure S2:**
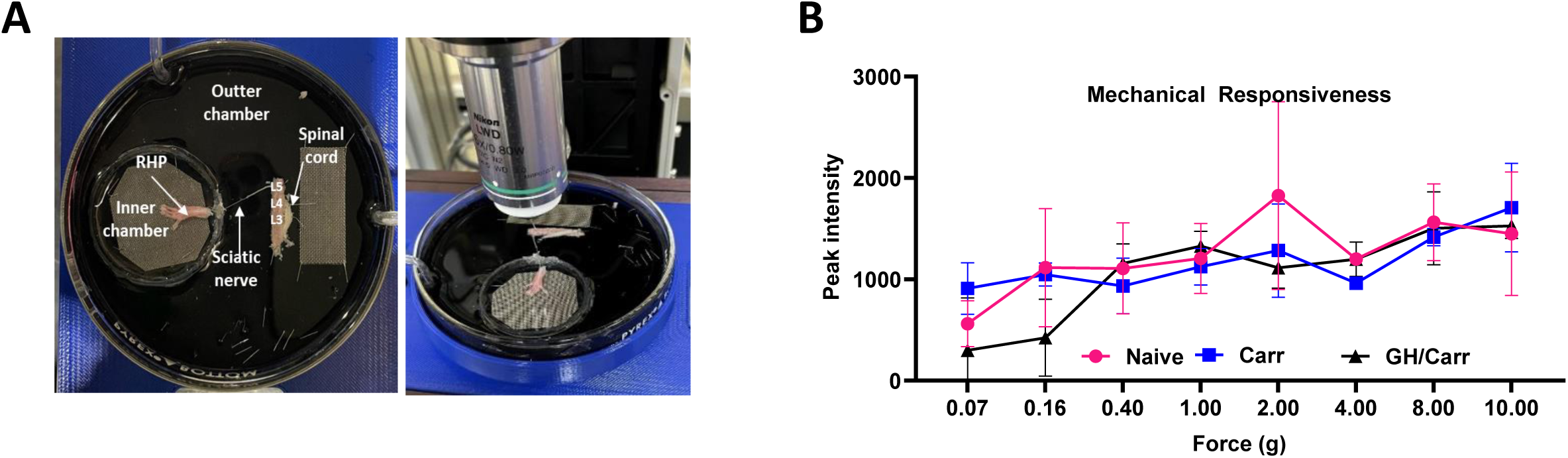
Representative *ex vivo* setup and mechanical responsiveness over increasing forces. **A.** Representation of the GCaMP *ex vivo* preparation and visualization using an upright confocal microscope. **B.** Peak fluorescence intensity to mechanical stimulation of the receptive fields at varying forces in naïve, carrageenan and GH/Carr groups.

**Figure S3:**
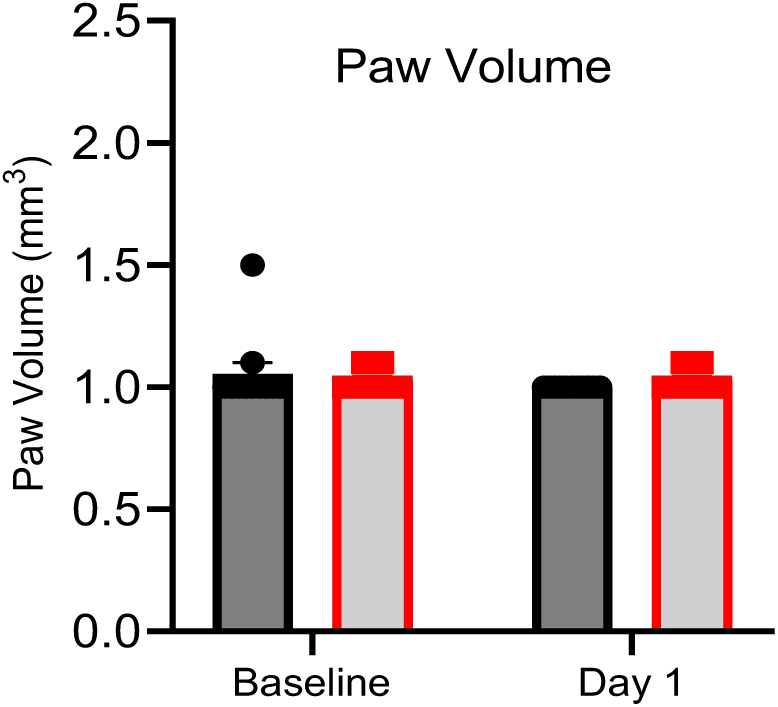
Contralateral paw volume in mice with carrageenan-induced muscle inflammation. Paw volume was not altered after muscle inflammation in the contralateral paw at day 1 when compared to baseline and naïve/sham groups. 1-way ANOVA.

**Figure S4:**
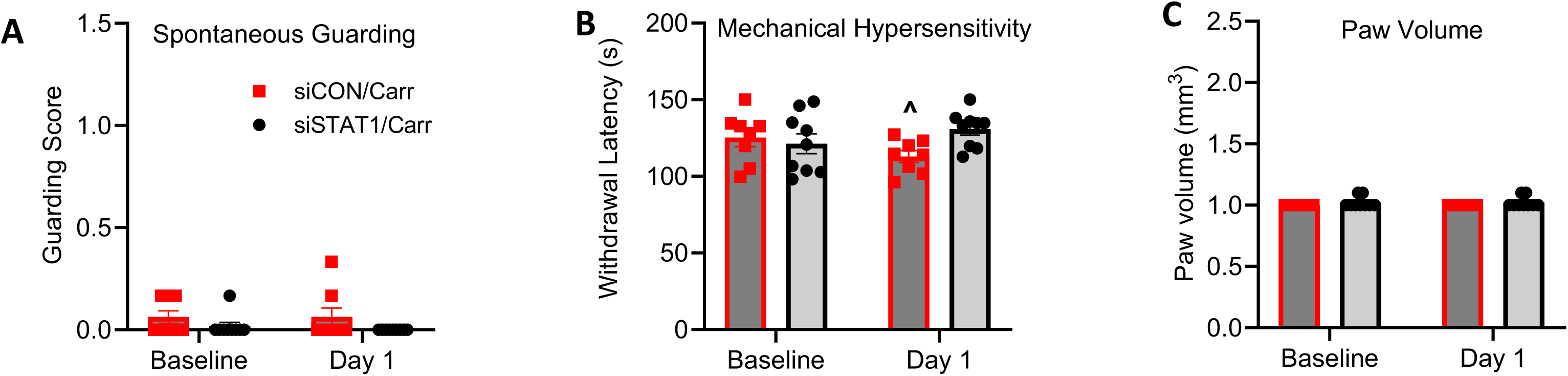
Nerve targeted MCP1 knockdown in sensory neurons alleviated contralateral hypersensitivity in neonatal mice. **A.** There are no observable changes in the guarding score in the contralateral limb of inflamed mice with siCON or siSTAT1 injection with carrageenan. **B.** However, contralateral muscle withdrawal thresholds in the same animals with siCON/Carr injections were slightly but significantly reduced compared with siSTAT1/Carr group (^p<0.05 vs siSTAT1/Carr) at day one. Two-way RM ANOVA followed by Tukey’s post hoc multiple comparison test. **C.** No changes in contralateral paw volume were observed in siRNA injected mice with muscle inflammation. n = 8-12/ group. Mean ± SEM.

**Figure S5:**
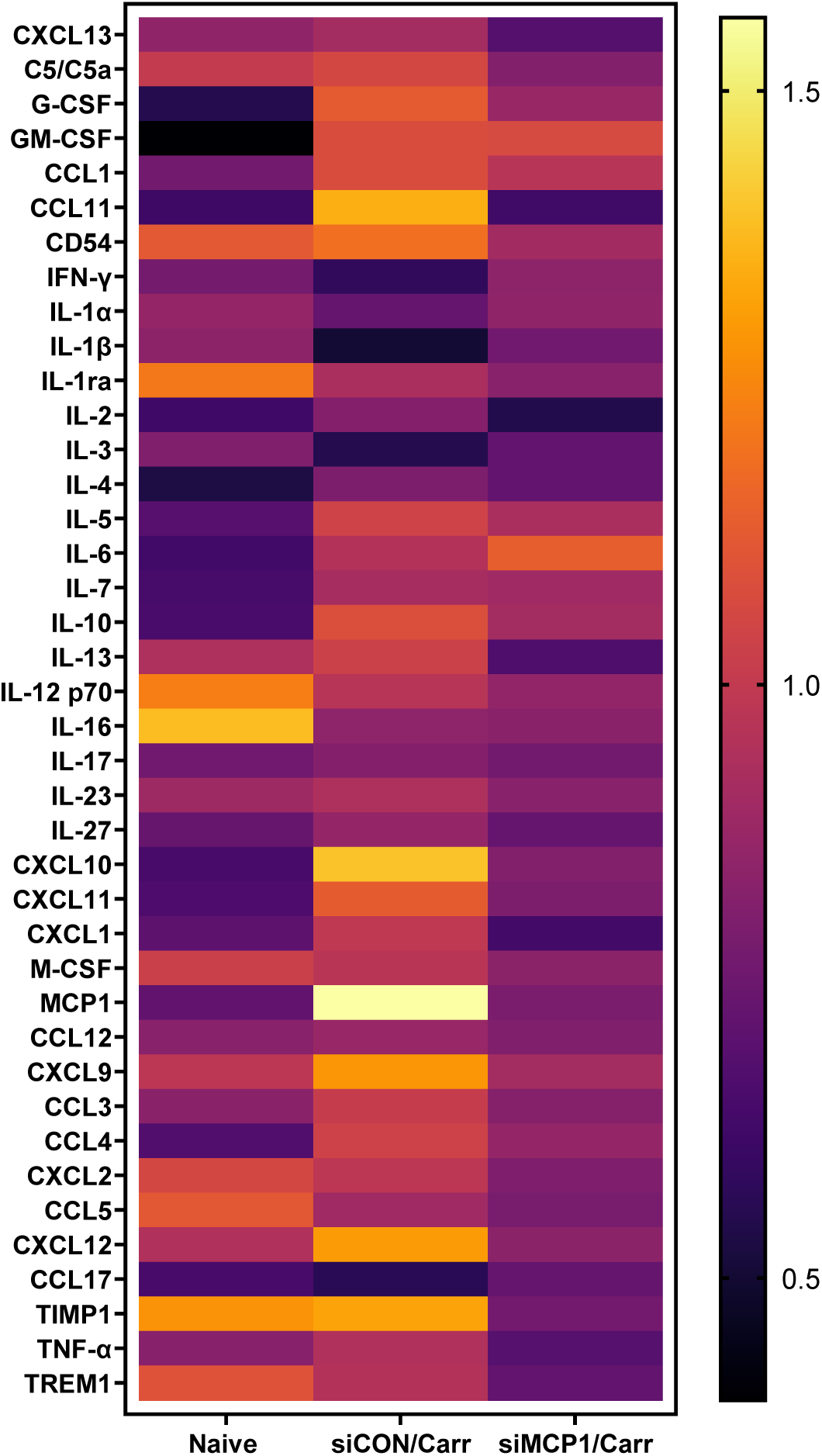
Representation of pro- and anti-inflammatory cytokines and chemokines expressed after muscle injury. Proteome Profiler Mouse Cytokine-Chemokine arrays were used to analyze factors in mice subjected to following inflammatory injury with siCON or siMCP1 injection and compared to naive. Average data from our replicates per group were represented via heat map.

**Table S1:** STAT1 is elevated in sensory neurons after cutaneous inflammatory injury. Data shown as a percent change from naive (p<0.05 vs. naive; n=3–6/group, One-way ANOVA with Tukey’s post hoc test).

| <u>% Change in DRG mRNA</u> |  |
| --- | --- |
| <u>Gene</u> | <u>1% Carr</u> |
| ELK1 | -10 +/- 20% |
| ELK3 | -35 +/- 23% |
| ELK4 | -26 +/- 22% |
| STAT1 | 278 +/- 20%* |
| STAT3 | -5 +/- 19% |
| STAT5a | -7 +/- 12% |
| STAT5b | -19 +/- 7% |
| SRF | -29 +/- 19% |

**Table S2:**
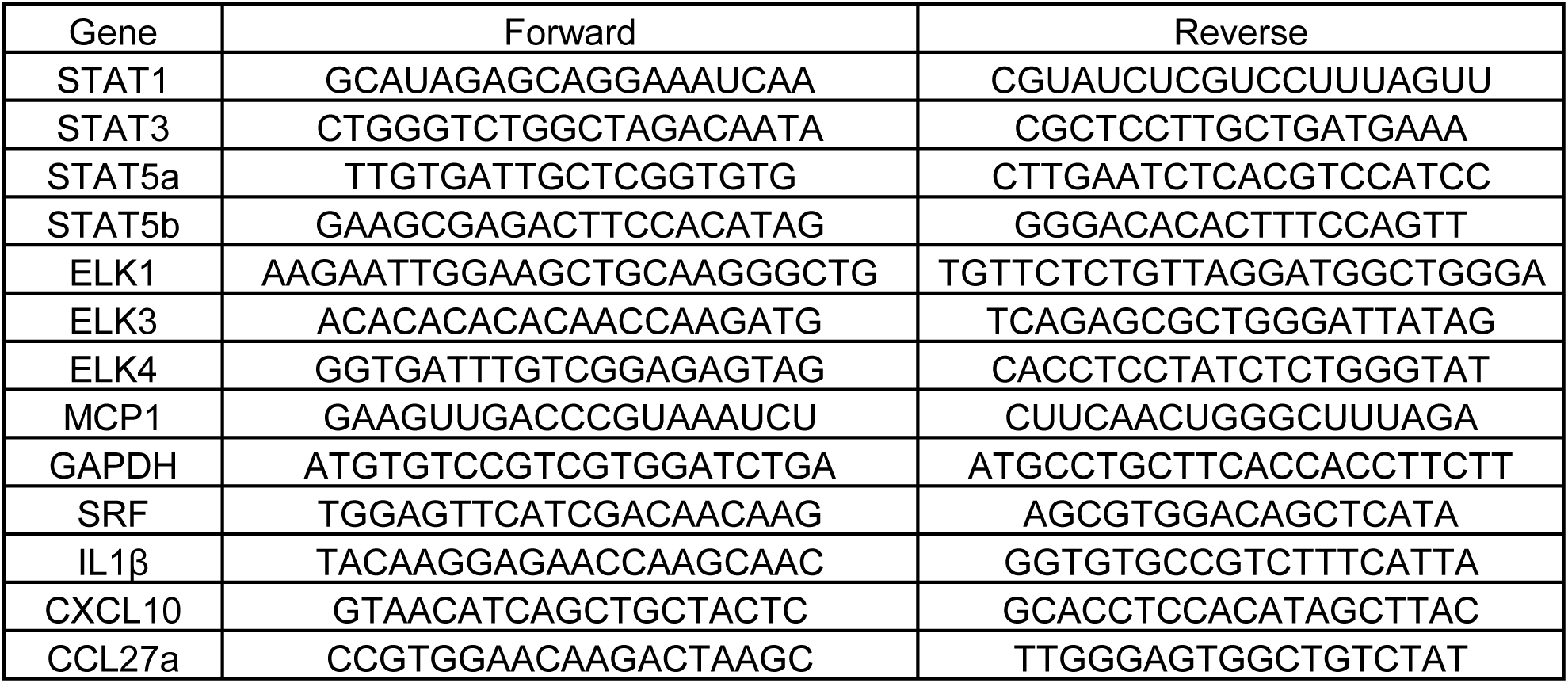
Primer sequences used for real time PCR in the current report.

